# Source-space precision charts for lifespan EEG connectomics

**DOI:** 10.64898/2026.06.19.732815

**Authors:** Yu Jin, Ronaldo Garcia Reyes, Ying Wang, Min Li, Maria L. Bringas-Vega, Pedro A. Valdes-Sosa

## Abstract

Source-space electroencephalography (EEG) connectomics aims to estimate interactions among cortical generators from sensor cross-spectra that are mixed by the head and lead field. This task is difficult because marginal source covariance or coherence can retain leakage, common drive and indirect mediation, whereas developmental mapping requires conditional interactions that can be estimated repeatedly across large cohorts. We developed JSPACE (Joint Source-space Precision And Cross-spectral Estimation), a frequency-domain inverse framework for estimating multi-frequency source precision matrices from scalp cross-spectra. JSPACE couples posterior source cross-spectral estimation with standardized precision fitting, sparse frequency-smooth anatomical regularization, stochastic active-set optimization and post-selection refitting. In simulations, its advantage was target-specific: JSPACE reduced coherence inflation and achieved the lowest imaginary-coherence and peak-frequency errors in a forward neural-mass benchmark. When the ground-truth precision matrix was known, it achieved the highest exact, edge-collapsed and leakage-aware support recovery. We applied JSPACE to HarMNqEEG cross-spectral data from 1,935 participants aged 5.17–97.00 years, spanning 47 frequency bins and 360 cortical parcels. Affine-invariant Karcher tangent harmonization reconstructed subject-level estimates into a lifespan atlas of 360 diagonal and 64,620 source-pair age-frequency surfaces. The atlas revealed a continuous off-diagonal morphology landscape, in which age direction, frequency preference and interaction strength varied as overlapping axes rather than discrete edge classes. In contrast, diagonal precision surfaces shared a conserved alpha-trough morphology across parcels. Representative real-precision pathways captured posterior parietal, sensorimotor-parietal, frontopolar and visual-parietal motifs. Delta-band gradients were moderately aligned with the sensorimotor-association (S-A) organization of cortex from childhood through late adulthood, with a candidate oldest-old deviation in the sparsest age range. JSPACE provides a scalable framework for frequency-resolved source-precision charting in lifespan EEG.

## 1 Introduction

Resting EEG and MEG resolve neuronal activity at millisecond time scales and are therefore well suited to frequency-resolved connectomics. However, the signals recorded at the scalp do not correspond one-to-one to cortical generators. Under the forward model, each sensor cross-spectrum is a mixture of source cross-spectra projected through the head and lead field [1, 2]. As a result, field spread and source leakage can make generators appear associated even when the apparent association is driven by spatial mixing, common drive or indirect pathways [3–5]. For electrophysiological connectomics, the inverse problem is therefore not only to localize oscillatory activity. It is also to estimate source-domain second-order structure in a form that remains interpretable after these confounds have been considered.

Classical and Bayesian source imaging methods provided the foundation for estimating local cortical activity from M/EEG, including minimum-norm, distributed tomographic and prior-constrained source solutions [6–8]. Frequency-domain extensions such as dynamic imaging of coherent sources made it possible to study oscillatory source power and coherence within an inverse framework [9]. In parallel, connectivity measures and corrections were developed to reduce sensitivity to spatial leakage, including the imaginary part of coherency, multivariate orthogonalization and explicit cautions about source-space ghost interactions [10–12]. These strategies address important sources of bias, but they do not by themselves define a statistical model of latent source interactions. Leakage-robust measures can lose sensitivity to near-zero-lag physiological coupling, whereas two-stage workflows can carry residual source-estimation errors into downstream pairwise connectivity measures [13]. Source connectivity therefore remains an inverse statistical problem rather than a post-processing problem.

This shifts the modeling target from source amplitudes alone to source cross-spectral statistics. Cross-spectral electrophysiological source imaging formalizes the estimation of latent source cross-spectra under the lead-field model, and spectral structured sparse Bayesian learning has shown that structured priors can reduce distortions in this high-dimensional inverse problem [14]. Cross-spectra and covariance matrices, however, describe marginal association. They do not distinguish a direct pairwise association from dependence induced through the rest of the modeled network. Precision matrices, the inverses of covariance or cross-spectral matrices, provide the corresponding conditional-dependence representation under Gaussian spectral models. Intuitively, an off-diagonal precision entry asks whether two sources remain statistically coupled after the activity of all other modeled sources has been taken into account. A zero off-diagonal entry corresponds to conditional independence in the Gaussian model, whereas covariance or coherence can remain nonzero because of common drive, indirect mediation or spatial mixing. Sparse inverse covariance estimation and joint graphical models show how regularization can recover interpretable graphs and borrow strength across related conditions [15, 16]. For M/EEG, hidden Gaussian graphical spectral models have already brought this idea into source-space inverse modeling through Hermitian graphical regularization [17].

Source-space precision estimation is not obtained by simply applying a graphical lasso after source reconstruction. The quantity available to the precision model is a posterior source moment shaped by the lead field, the residual sensor model and the regularization used in the inverse step. If this moment is passed directly to a precision fit, local spectral power differences can dominate the graph and obscure weaker conditional dependencies. Fitting each frequency independently introduces a second instability, because neighboring spectral bins carry related physiological and sampling information. Anatomical connectivity provides another constraint, but it should act as a soft source of regularization rather than imposing a hard support constraint a priori. These considerations lead to a coupled formulation in which source-moment estimation and precision fitting are linked, and in which standardization, frequency sharing and anatomical weighting are built into the estimator.

The need to estimate comparable source-space conditional dependence becomes more demanding in lifespan and normative EEG. A population atlas requires the same inverse estimator to be applied across many subjects, sites and ages. Its outputs must also remain comparable after harmonization. qEEGt addressed this population problem for EEG source spectra through age-corrected normative statistical parametric mapping, and HarMNqEEG extended the framework to harmonized multinational EEG cross-spectral tensors [18, 19]. Recent pipelines and models, including CiftiStorm and *ξ*-*α*NET, further show that source-space EEG can be organized at large scale and connected to structural priors, lifespan modeling and reproducible workflows [20, 21]. The unresolved question is whether this population neuroinformatics framework can be extended from local spectra or specific generative-model parameters to frequency-resolved source conditional-dependence charts.

Here we introduce JSPACE (Joint Source-space Precision And Cross-spectral Estimation) to address this question. JSPACE couples posterior source cross-spectral estimation with multi-frequency Hermitian precision fitting. Its central modeling step is to standardize posterior source second moments before precision estimation, so that conditional-dependence structure is not driven primarily by local power differences. The precision model then combines sparse off-diagonal regularization, smoothness across neighboring frequencies and soft anatomical weighting, and the fitted precision is returned to the source scale for the next inverse update. We evaluate JSPACE in simulations that separate source-spectral recovery from precision-matrix recovery, and then apply it to HarMNqEEG data to build a harmonized lifespan atlas of reconstructed source-precision metrics. Together, the simulations and HarMNqEEG application evaluate JSPACE as an estimator of frequency-resolved conditional dependence among cortical sources and as a framework for constructing cohort-scale lifespan source-precision charts.

## 2 Materials and Methods

The Methods define the statistical objects needed to interpret the validation and developmental results. We first specify the empirical cross-spectral input and source space, because every subsequent precision estimate is conditional on this forward model. We then describe the subject-level JSPACE estimator, the two benchmark settings used to separate source-spectral recovery from precision recovery and the Karcher tangent-space procedure used to turn individual precision estimates into developmental charts. The main text gives only the summary operations needed to read the atlas figures; detailed equations, solver rules, numerical constants, screening thresholds and diagnostic tables are kept in the appendices.

### 2.1 Cohort, source space and analysis scope

The empirical application used HarMNqEEG cross-spectral data from a multisite lifespan EEG resource [19]. This dataset provides harmonized scalp spectral summaries rather than raw EEG for the present analysis, so the modeling begins from subject- and frequency-specific scalp cross-spectra. The present study used de-identified derived cross-spectral summaries and did not access identifiable participant information or raw EEG recordings; ethical approvals, consent procedures and recruitment criteria for the contributing datasets are described in the HarMNqEEG resource paper [19]. Seventeen of 1952 participant-level records lacked usable age information and were excluded before model fitting, yielding *N* = 1935 participants aged 5.17–97.00 years. No additional participants were removed during JSPACE estimation. The source model used 360 Human Connectome Project multimodal parcellation (HCP-MMP1) parcels on a fixed parcel lead field [22]. Cohort geography, site sample sizes, reported sex/gender availability and age coverage are summarized in Figure 1; site-level characteristics are provided in Appendix B.

**Figure 1.**
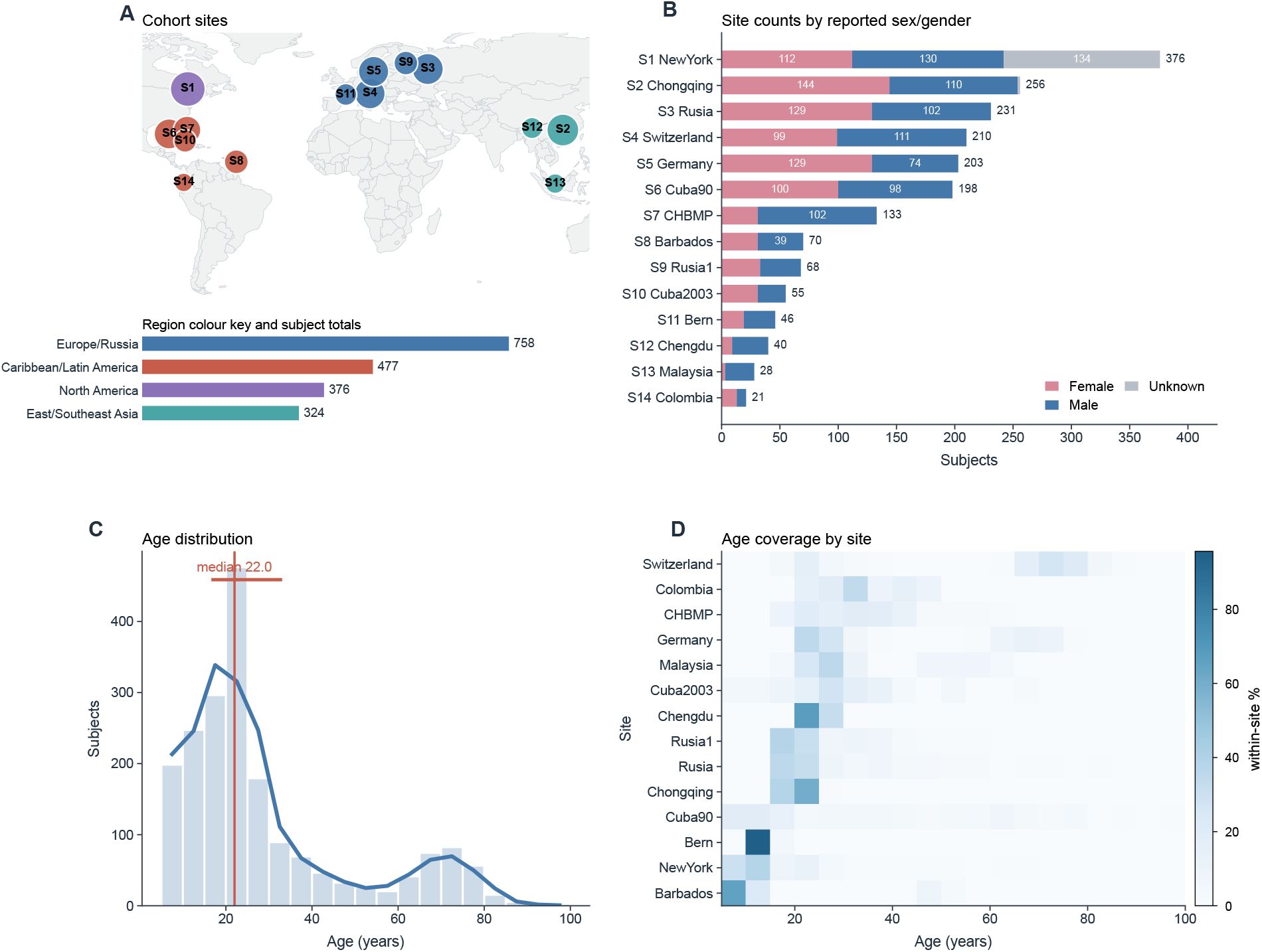
HarMNqEEG cohort geography, site composition and age coverage. **A**, recording-site locations; marker area scales with retained subjects and marker color denotes region, with regional totals shown below the map. **B**, site-level sample sizes by stacked reported sex/gender counts. **C**, overall age distribution with the cohort median. **D**, age coverage across recording sites.

**Table 1.**
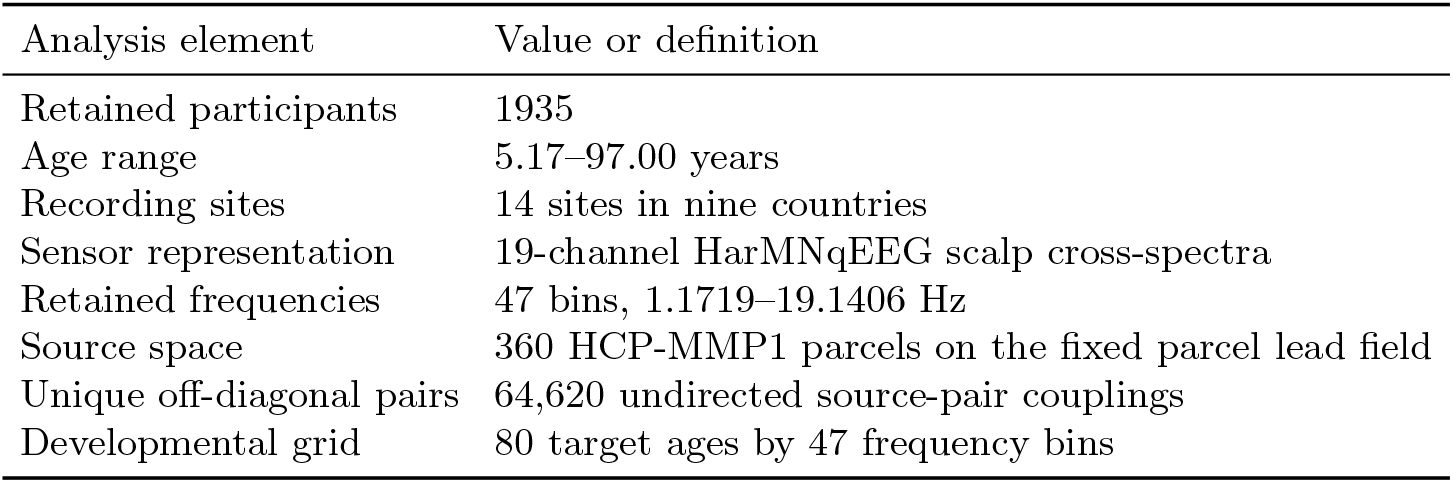
Main empirical analysis scope. Detailed site-level cohort characteristics are reported in Appendix Table B.1.

For subject *i* and frequency *f*, JSPACE treats the HarMNqEEG scalp cross-spectrum 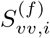 as a fixed empirical input. Source inference used a parcel lead field 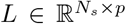, with *N*_*s*_ = 19 sensors and *p* = 360 parcels, obtained from the upstream CiftiStorm–Brainstorm/HCP-MMP1 workflow. The source cross-spectral model was

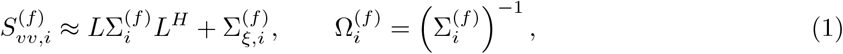

where 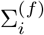 is the latent source covariance, 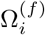 is the source precision matrix and 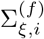 is a diagonal sensor residual covariance. The empirical inverse problem is strongly underdetermined because the source dimension is 360 and the sensor dimension is 19. The source-precision estimates are therefore not treated as direct cortical observations; they are conditional estimates defined by the forward model, parcellation, residual model, regularization and post-selection procedure.

### 2.2 JSPACE source-precision estimator

The source model in Eq. 1 leaves two coupled quantities unknown: the posterior source second moment and the precision matrix that regularizes it. A post-hoc inversion of a source covariance would separate these two quantities, although each depends on the other in a low-sensor, high-source inverse problem. JSPACE therefore alternates a posterior source-moment update with a standardized multi-frequency precision update (Fig. 2). At outer iteration *q*, the posterior step produces a source second moment 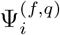. Before precision fitting, this moment is standardized so that marginal source scale is not allowed to dominate the off-diagonal conditional-dependence structure:

**Figure 2.**
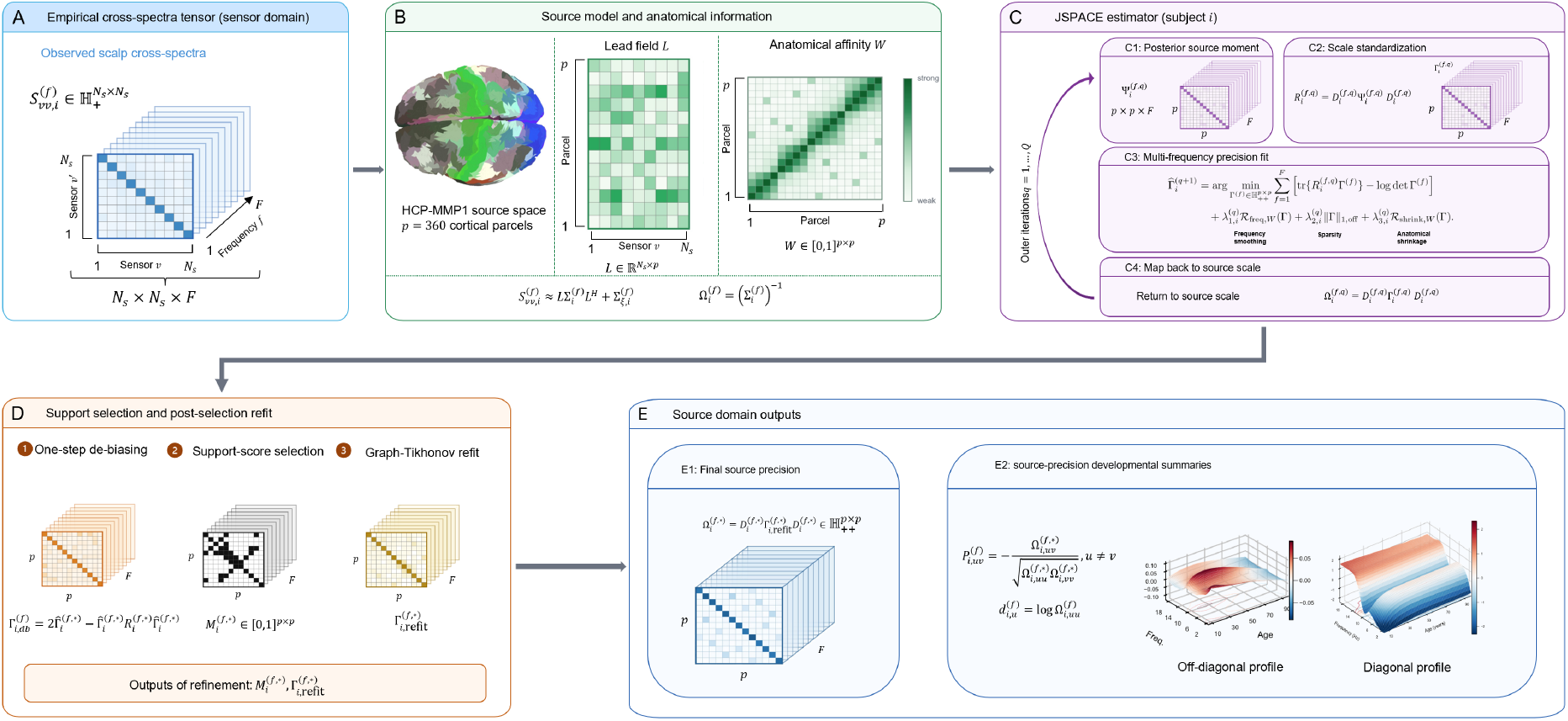
JSPACE source-precision estimation pipeline. **A**, Subject- and frequency-specific scalp cross-spectral tensors provide the sensor-domain input. **B**, The inverse model combines the HCP-MMP1 parcel source space, a 19-sensor lead field and an anatomical affinity matrix used as a soft regularization weight. **C**, For each subject, JSPACE alternates posterior source-moment estimation, scale standardization and multi-frequency standardized precision fitting, with effective penalties induced by the subject-level profile *η*_*i*_, before mapping the standardized estimate back to source scale. **D**, Post-selection processing uses one-step de-biasing to form a dense screening statistic, trace-minus-log-determinant support-score selection to define the support mask and a frequency-smoothed graph-Tikhonov refit on the selected support. **E**, The final source-domain precision tensor and derived normalized precision summaries provide the inputs for diagonal and off-diagonal developmental surface analyses.

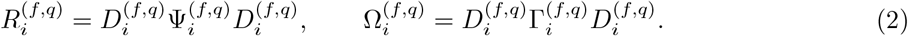

Here 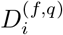 is a diagonal standardization matrix, 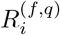 is the standardized posterior moment and 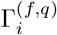 is the standardized precision matrix. This decomposition lets the likelihood be fitted on a scale-controlled object while still returning the final estimate to the original source scale.

At each outer iteration, the standardized precision sequence is fit jointly across the *F* = 47 retained frequencies. The frequency dimension is modeled jointly because adjacent spectral bins are expected to share conditional-dependence structure, whereas fitting each bin independently would amplify sampling noise in a 360-parcel graph. A fixed subject-level regularization profile *η*_*i*_ = (*η*_1,*i*_, *η*_2,*i*_, *η*_3,*i*_) is selected once for each subject. Its values induce iteration-specific effective penalties 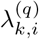 through scale factors computed from the current standardized posterior moment:

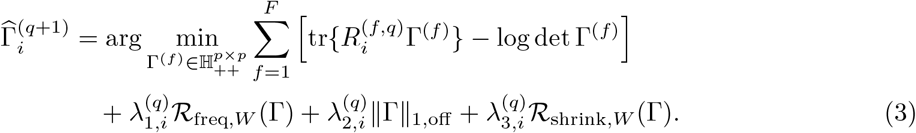

The likelihood term fits a Gaussian precision model to the standardized posterior moment. In this notation, 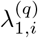 controls frequency smoothing, 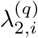 controls off-diagonal sparsity and 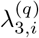 controls anatomically weighted shrinkage. The weighted shrinkage and frequency penalties use parcel-level anatomical affinity weights from the upstream structural prior as soft anatomical information, consistent with structurally informed approaches to functional and electrophysiological connectivity estimation [23,24]. This weighting shrinks anatomically weaker pairs more strongly and encourages smoother trajectories across neighboring frequencies. Thus *η*_*i*_ defines the subject-level penalty profile, whereas 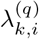 is the effective penalty applied at iteration *q*. Exact definitions of 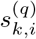, the mapping from *η*_*i*_ to 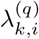, *ℛ*_freq,*W*_,ℛ_shrink,*W*_, hyperparameter search, stochastic optimization, one-step de-biased support screening and post-selection refitting are given in Appendix C and Appendix D.

Because the off-diagonal *ℓ*_1_ term shrinks conditional-dependence estimates toward zero, the penalized standardized precision was not used directly as the post-selection support statistic. After the final M-step, JSPACE formed a one-step de-biased precision statistic in standardized source space and used this statistic for Rayleigh support screening. The selected support was then held fixed during post-selection refitting, so the reported source precision matrices are refitted estimates rather than raw thresholded de-biased matrices.

The final standardized estimate is mapped back to source scale before downstream analysis. Off-diagonal source precision is reported as a normalized precision coefficient,

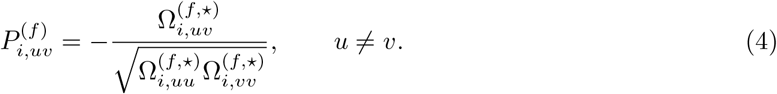

The real part, imaginary part, magnitude and phase of 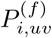 define distinct conditional-precision summaries. These summaries are not interchangeable with source power, coherence or directed connectivity; they are different projections of the same estimated source-domain precision object.

### 2.3 Simulation benchmarks

The validation strategy follows from the distinction between source-spectral reconstruction and precision estimation. A forward neural-mass benchmark asked whether JSPACE preserved source-spectral and phase-lagged structure after forward mixing through the same lead-field representation used in the empirical analysis. This benchmark is useful for source-domain spectral behavior, but it is not a precision-graph ground truth because the simulated source cross-spectrum can be low rank when the ensemble count is smaller than the number of parcels. A second reduced-dimensional benchmark therefore generated sparse Hermitian positive-definite precision matrices with smooth frequency-varying off-diagonal trajectories. That benchmark directly evaluated precision and partial-correlation errors, exact support recovery and leakage-aware spatial support recovery. Full simulation equations and numerical settings are given in Appendix E.

### 2.4 Karcher tangent-space developmental atlas

The developmental analysis is a downstream application of the final JSPACE precision matrices, not part of the subject-level estimator. Because these matrices are Hermitian positive-definite (HPD), their averaging and smoothing should respect matrix geometry rather than treating every entry as an unrelated scalar. We therefore used an affine-invariant Karcher reference for each frequency. Denoting the stabilized final precision matrix for subject *i* and frequency *f* by 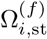, the tangent representation was

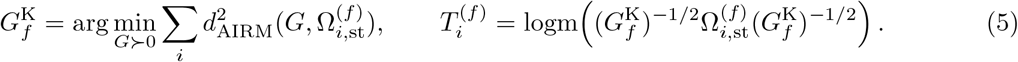

Here *d*_AIRM_ is the affine-invariant Riemannian distance on HPD matrices. This geometry is used because HPD matrices do not form a flat Euclidean space: entrywise averaging can distort eigenvalues and does not respect the multiplicative scale of covariance or precision matrices. The affine-invariant metric compares matrices after whitening by the reference matrix, so tangent coordinates describe relative departures from the Karcher mean rather than raw elementwise differences. The Hermitian tangent matrix 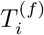 was then vectorized into coordinates *τ*_*ifh*_, giving 129,600 real components per frequency: 360 diagonal terms, 64,620 real off-diagonal terms and 64,620 imaginary off-diagonal terms.

Uneven age sampling and multisite acquisition can otherwise confound lifespan summaries of these tangent components [25, 26]. Each component was therefore harmonized in age-frequency space. Let *x*_*i*_ = log(*a*_*i*_), *z*_*f*_ = log(*ν*_*f*_) and *s*_*i*_ denote the acquisition site. A global location-scale surface produced standardized residuals, and site-specific location-scale surfaces were estimated on the standardized scale:

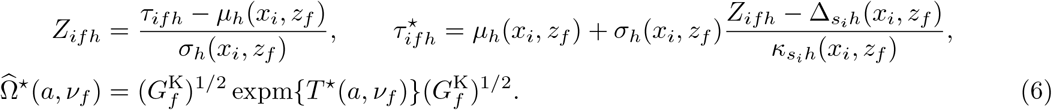

Here *τ*^⋆^(*a, ν*_*f*_) is the grid-level local-polynomial mean of the harmonized tangent coordinates, and *T* ^⋆^(*a, ν*_*f*_) is the Hermitian matrix obtained by unvectorizing that predicted coordinate vector. Biological interpretation is based on reconstructed harmonized precision metrics, including 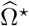 and 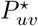, not on standardized residual maps.

Atlas-wide descriptive summaries were used to make the reconstructed metric bank readable at manuscript scale. For off-diagonal surfaces, normalized surface-shape features were embedded with uniform manifold approximation and projection (UMAP) as a visualization of the continuous morphology landscape. Continuous feature scores for age direction, alpha-axis structure, low-frequency dominance, age-frequency interaction and midlife curvature were then used to annotate that landscape and to select representative surfaces. For the diagonal layer, parcel surfaces were standardized before averaging and similarity analysis, allowing the conserved frequency profile and parcel-level alpha-trough depth to be summarized across the 360 parcels. For selected real-precision pathways, the full reconstructed age-frequency surface was shown together with frequency and age slices.

For the delta-band hierarchy analysis, age-specific parcel gradients were computed from delta-band |*P*^⋆^| profiles by spectral embedding of parcel-wise precision profiles. The gradient sign was oriented to an external sensorimotor-association reference projected to HCP-MMP1 parcel space. Alignment with this reference was quantified using Spearman correlation and spatial spin-null testing, while pairwise map similarity and multidimensional scaling summarized age-specific reorganization. Full screening rules, projection details, feature definitions and phase-specific caveats are provided in Appendix F.

### 2.5 Computational implementation and reproducibility

All subject-level JSPACE fits used fixed analysis settings after preliminary subject-level hyperparameter optimization. JSPACE fitting and hyperparameter search were implemented in MATLAB R2026a (MathWorks, Natick, MA, USA), with stochastic search controlled by the fixed seeds and common-random-number settings reported in Appendix D. The developmental atlas processed all 1935 retained participants, 47 frequency bins, 360 parcels and 129,600 tangent components per frequency before reconstruction of the metric bank. Manuscript figure generation and source-data export used Python 3 with NumPy, SciPy, pandas, h5py and Matplotlib; exact package versions and execution scripts are provided with the code repository. Computations for the current reproducibility build were performed on a 64-bit Windows workstation with an Intel Core i9-14900KF CPU, 32 logical processors and 128 GB RAM. Data and code access conditions are stated in the Data and Code Availability section.

## 3 Results

### 3.1 Forward simulations separate source-spectral reconstruction from precision estimation

We first evaluated JSPACE in a forward neural-mass benchmark that emphasized source-spectral reconstruction after sensor mixing. This benchmark was necessary because the comparator workflows, *ξ*-*α*NET, MNE+FOOOF and eLORETA+*ξ*-*α*, primarily estimate source spectra or covariance summaries, whereas JSPACE directly estimates regularized source precision matrices. This benchmark therefore separated source-spectral recovery from the precision-graph target evaluated below.

The benchmark passed time-domain source dynamics through the same lead-field representation used in the empirical analysis. In this setting, *ξ*-*α*NET had the lowest source-power, log-power and source-covariance errors (Table 2). JSPACE had larger power and covariance errors than *ξ*-*α*NET, so the forward benchmark did not identify it as the lowest-error estimator of local spectral amplitude. Its behavior differed on cross-spectral summaries: JSPACE avoided the large coherence errors observed for MNE+FOOOF and eLORETA+*ξ*-*α*, and it had the lowest imaginary-coherence and peak-frequency errors. The forward benchmark therefore constrained the interpretation of the method before the developmental atlas was analyzed: JSPACE preserved frequency and phase-lagged cross-spectral structure after forward mixing, but its main target was not source-power minimization.

**Table 2.**
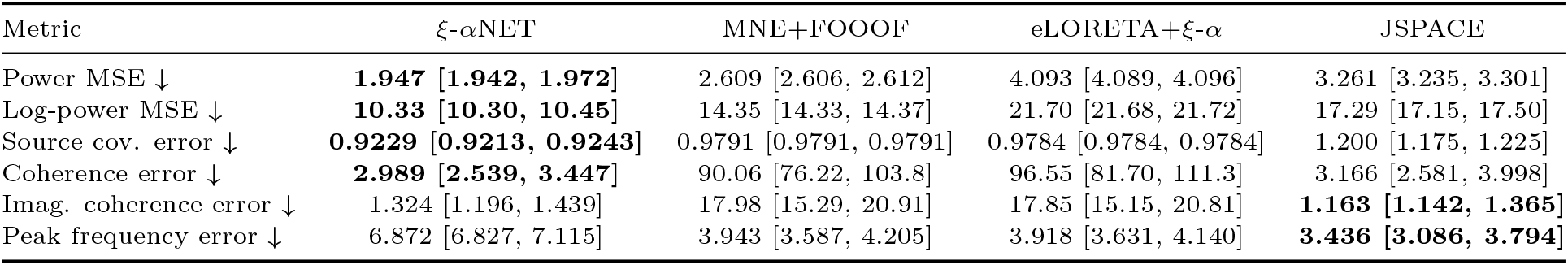
Forward benchmark: covariance and source-spectral recovery. Values are median [IQR] across 50 simulation replicates. Arrows indicate the preferred direction, and bold marks the best median.

### 3.2 JSPACE improves recovery when source precision is the target

The second benchmark matched the simulated ground truth to the estimand of JSPACE. Sparse Hermitian positive-definite precision matrices were generated with smooth frequency-varying off-diagonal trajectories and then projected through the forward model. This setting directly tested precision and normalized partial-correlation error, exact graph support recovery and leakage-aware spatial support recovery.

The target-matched benchmark showed that the advantage of JSPACE was concentrated in support recovery and spatially tolerant edge localization (Figure 3). Exact edge-by-frequency recovery was difficult for all methods, but JSPACE had the highest median exact support F1 (0.1226 [0.1168, 0.1261]) and edge-collapsed F1 (0.1496 [0.1438, 0.1567]). The advantage became clearer when evaluation allowed for the spatial ambiguity expected in EEG source imaging. JSPACE achieved the highest lead-field tolerant recall for both top-3 neighborhoods (0.7356 [0.6883, 0.7566]) and top-5 neighborhoods (0.9098 [0.8914, 0.9212]), and the highest PSF/CTF tolerant top-5 recall (0.9658 [0.9208, 0.9864]). It also had the smallest true-to-estimated edge distance (0.1819 [0.1728, 0.1921]) and symmetric edge distance (0.2053 [0.1993, 0.2099]).

**Figure 3.**
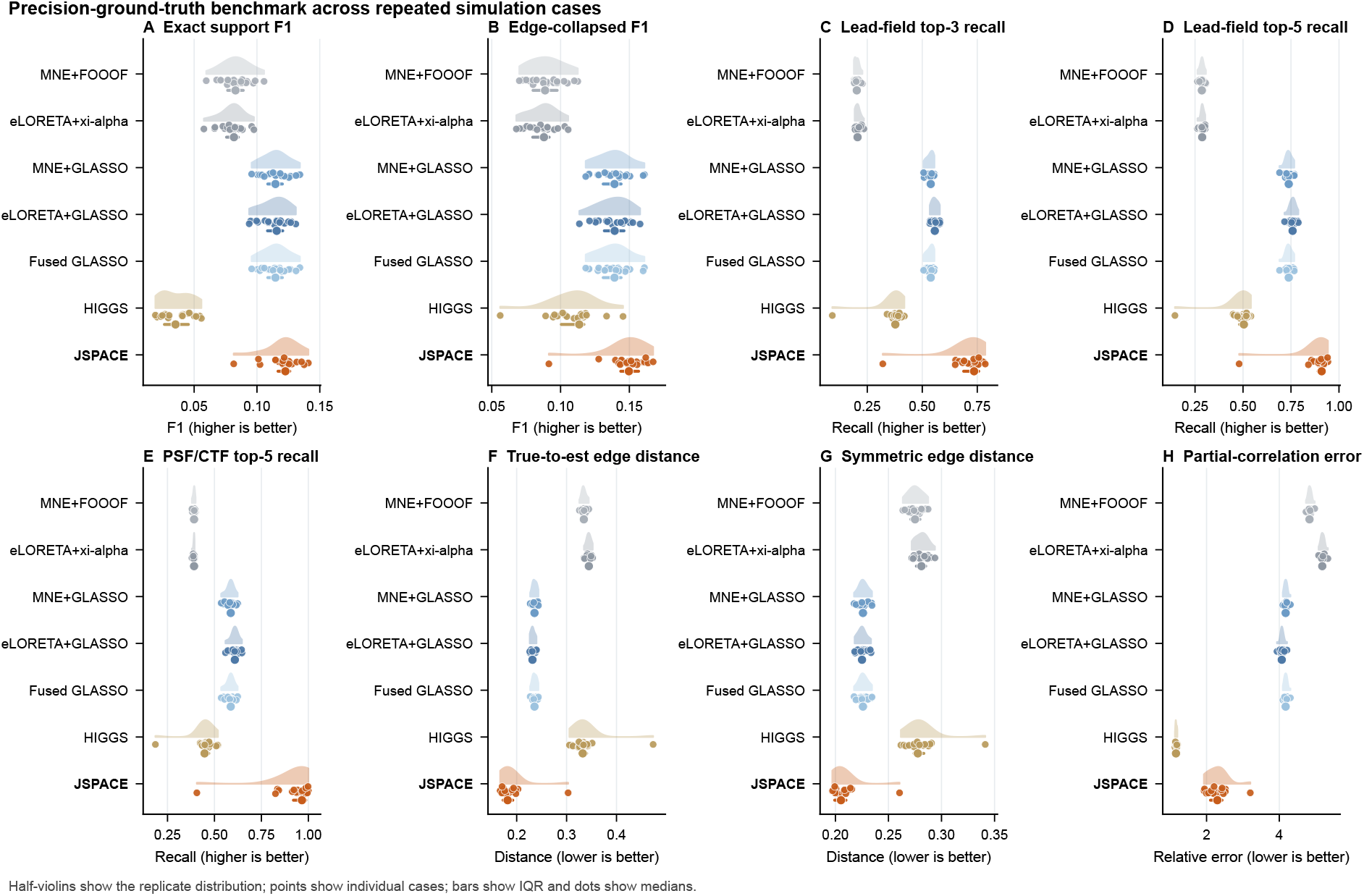
Precision-ground-truth benchmark across repeated simulation cases. Half-violins show the distribution across 20 precision benchmark replicates, points show individual cases, horizontal bars show the interquartile range and dots show medians. The display includes source-workflow baselines, post-hoc graphical-lasso baselines, HIGGS and JSPACE. **A**, Exact edge-by-frequency support F1. **B**, Edge-collapsed support F1 after collapsing support across frequencies. **C**,**D**, Lead-field tolerant edge recall using the top-3 and top-5 lead-field-confusable parcel neighborhoods. **E**, PSF/CTF tolerant top-5 edge recall using neighborhoods derived from point-spread/cross-talk structure. **F**,**G**, Region-localization distance from true to estimated edges and the corresponding symmetric edge distance. **H**, Relative Frobenius error of normalized partial-correlation matrices. Higher values are better in panels A–E; lower values are better in panels F–H.

This pattern separated graph-support localization from scalar partial-correlation amplitude recovery. JSPACE remained below the post-hoc graphical-lasso and source-workflow baselines on partial-correlation error (2.297 [2.108, 2.425]), but it was not the lowest-error method for that scalar target. Thus, the precision benchmark did not identify JSPACE as uniformly best on every numerical summary. Instead, it showed that JSPACE was strongest for recovering the location of sparse conditional-dependence structure under forward-model ambiguity, which is the validation target most closely matched to the downstream lifespan precision atlas.

### 3.3 Off-diagonal precision surfaces form a continuous morphology landscape

After the simulations, we applied JSPACE to the HarMNqEEG cohort and reconstructed a harmonized lifespan precision atlas from the final subject-level matrices. The atlas covered all 1935 retained participants, 47 frequency bins, 360 HCP-MMP1 parcels and 64,620 unique undirected source-pair couplings. We first asked whether the off-diagonal precision surfaces could be reduced to a small set of discrete classes, or whether they varied along continuous age-frequency axes.

The off-diagonal surfaces formed a continuous morphology landscape rather than a set of sharply separated classes (Figure 4A). In this display, each source pair was represented by normalized surfaceshape features and embedded with UMAP for visualization. The embedding was not used as an inferential clustering result. Instead, it provided a map of how reconstructed age-frequency surfaces varied across the atlas. Representative anchors showed several interpretable surface morphologies, including late increase, late decrease, alpha trough, alpha ridge, low-frequency dominance and strong age-frequency interaction (Figure 4C).

**Figure 4.**
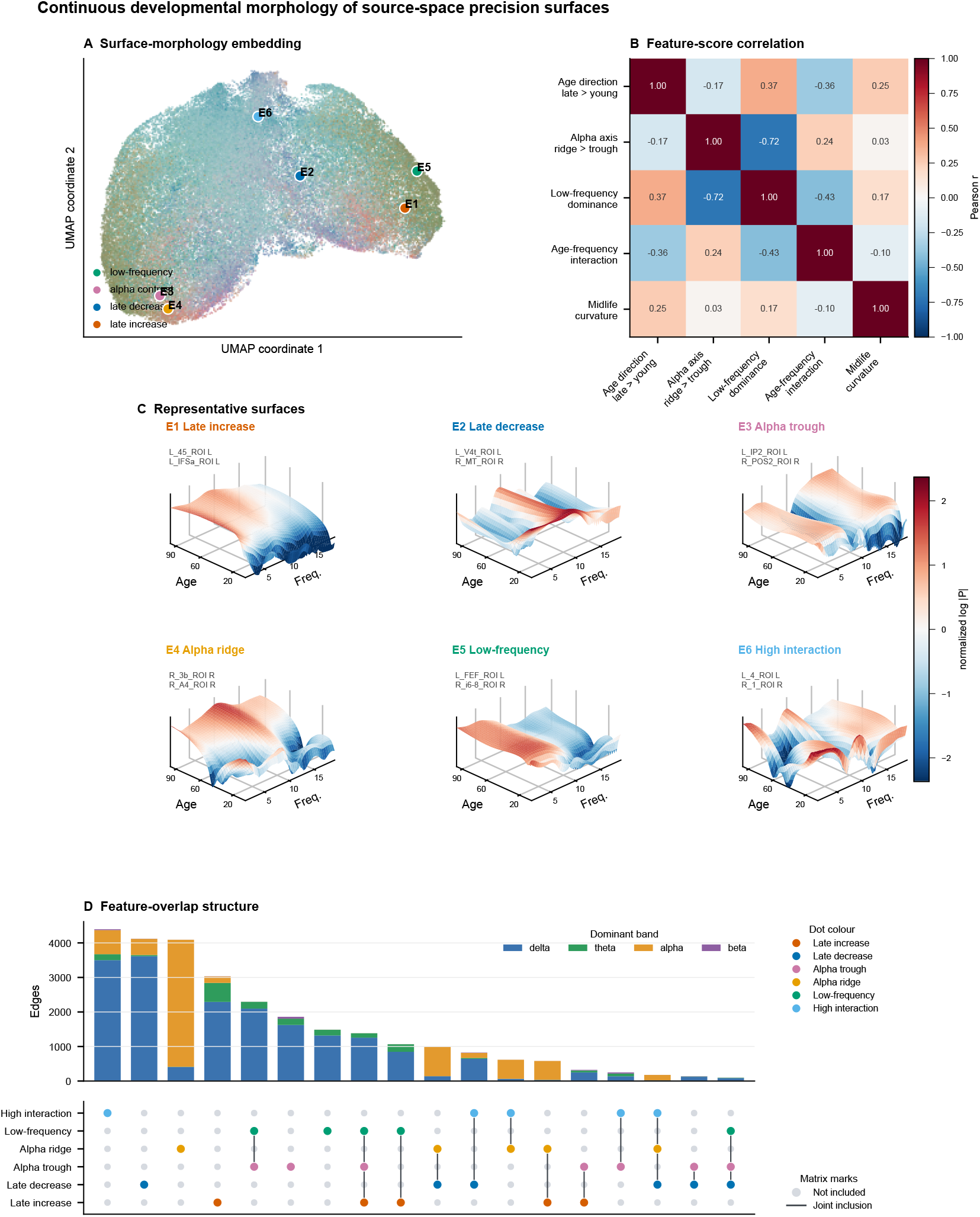
Continuous morphology landscape of reconstructed off-diagonal source-precision surfaces. **A**, UMAP visualization of normalized off-diagonal age-frequency surface morphology across 64,620 source pairs. Colors blend continuous feature scores, and labeled anchors mark representative surfaces shown in panel C. **B**, Correlation structure among continuous feature scores, showing overlap and opposition among age direction, alpha-axis structure, low-frequency dominance, age-frequency interaction and midlife curvature. **C**, Representative reconstructed normalized log-magnitude precision surfaces selected from feature extremes. The examples illustrate late increase, late decrease, alpha trough, alpha ridge, low-frequency dominance and high age-frequency interaction. **D**, Descriptive feature-overlap summary for feature-extreme source pairs. Bars show the number of source pairs in each motif combination, colored by dominant frequency band. Filled dots indicate motifs included in each combination. The feature-extreme sets provide an atlas-level description of surface organization and were not used to define an edgewise inferential set.

The continuous feature scores were not independent dimensions. Alpha-axis structure was strongly opposed to low-frequency dominance (*r* = −0.72), whereas age direction was positively related to low-frequency dominance (*r* = 0.37) and negatively related to age-frequency interaction (*r* = −0.36; Figure 4B). These relationships indicate that the atlas is not well described by one-dimensional labels such as “developmental increase” or “alpha effect”. A surface could express age direction, spectral localization and age-frequency interaction to different degrees.

To summarize this overlap without treating the motifs as mutually exclusive biological classes, we examined feature-extreme source pairs for the main continuous axes (Figure 4D). This descriptive summary showed that many source pairs expressed more than one motif: 18,983 source pairs met one descriptive feature-extreme criterion, 7,212 met two, 1,775 met three and 10 met four, while 36,640 met none. The band composition of the largest intersections was dominated by delta and alpha contributions, consistent with the representative surfaces in panel C. Thus, the first atlas-level result is that off-diagonal source precision is organized as a continuous, overlapping age-frequency morphology landscape rather than as a set of isolated edge types.

### 3.4 Diagonal precision surfaces share a conserved alpha-trough architecture

The diagonal entries of the precision matrix showed a different form of organization from the offdiagonal coupling surfaces. Whereas off-diagonal source pairs occupied a continuous morphology landscape, diagonal log precision was dominated by a highly reproducible spectral architecture across cortical parcels (Figure 5). Because diagonal precision represents the local conditional scale of a source after accounting for the rest of the graph, we analyzed it separately from inter-parcel coupling.

**Figure 5.**
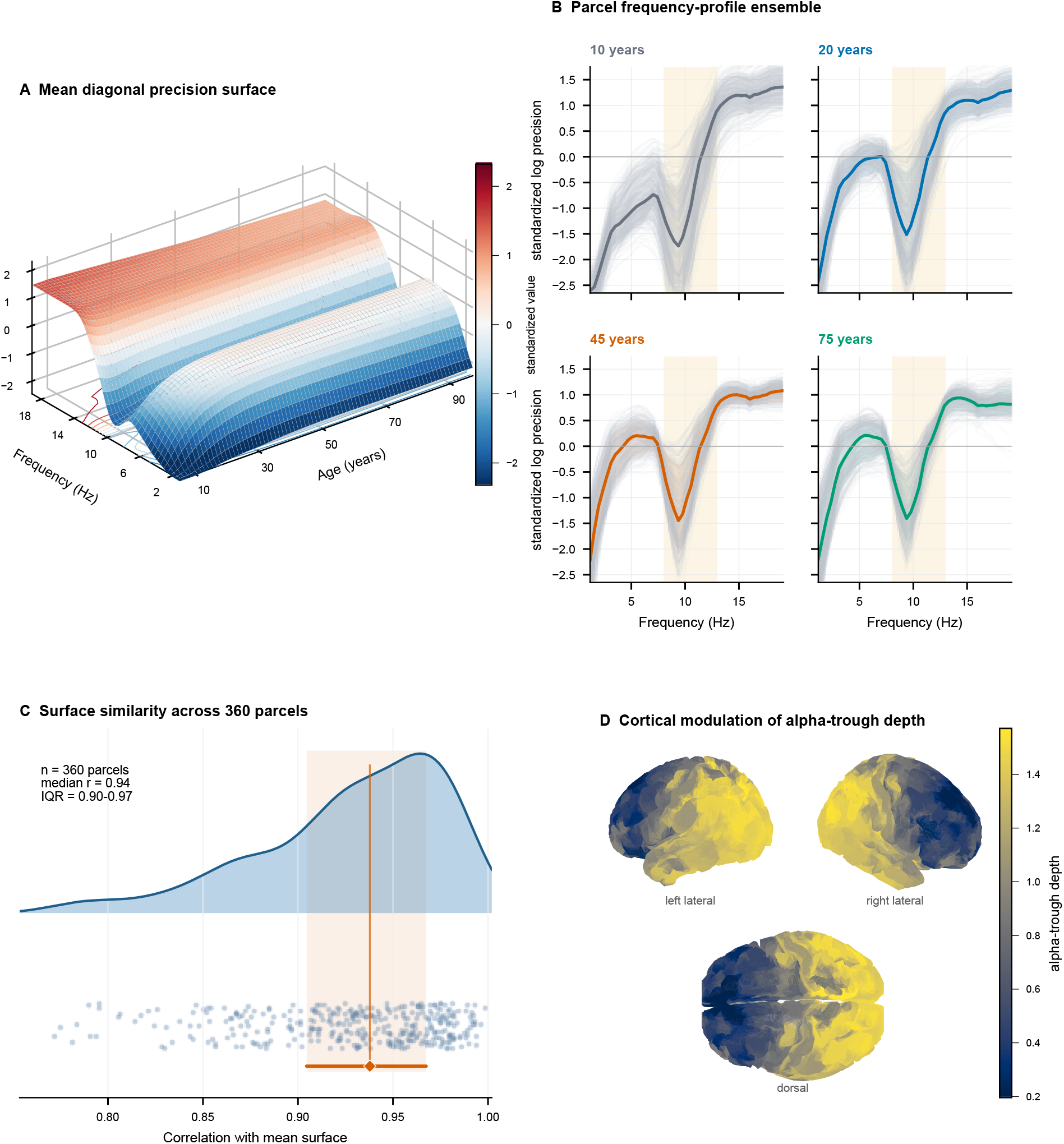
Conserved alpha-trough structure in diagonal conditional precision. **A**, Mean standardized diagonal log-precision surface across 360 HCP-MMP1 parcels. **B**, Parcel frequency-profile ensembles at representative ages. Gray lines show individual parcel profiles, colored lines show the parcel mean at each age, and the shaded interval marks the alpha range. **C**, Full-surface similarity between each parcel and the atlas mean diagonal surface, quantified by Pearson correlation after within-parcel standardization. The vertical line marks the median correlation, and the horizontal interval marks the interquartile range. **D**, Cortical map of alpha-trough depth, defined for each parcel as the mean of the theta and low-beta flanks minus the mean alpha-band value in the standardized diagonal surface.

The mean standardized diagonal surface showed a pronounced trough in the alpha range, embedded within a broader increase toward higher retained frequencies (Figure 5A). This pattern was not driven by a small subset of parcels. Across all 360 HCP-MMP1 parcels, each parcel surface remained strongly correlated with the atlas mean surface (median *r* = 0.938, IQR 0.905–0.967; minimum *r* = 0.772; Figure 5C). Frequency profiles sampled at representative ages showed the same alpha-centered depression across childhood, early adulthood, midlife and older adulthood (Figure 5B).

We quantified the depth of this alpha trough as the mean of the theta and low-beta flanks minus the mean alpha-band value within each standardized parcel surface. The trough depth was positive in every parcel (*n* = 360; minimum 0.088, median 1.096, maximum 1.664), indicating that the alpha depression was a conserved feature of local conditional precision rather than an averaging artifact. Its magnitude nevertheless varied across cortex (Figure 5D). The largest trough depths were concentrated in bilateral posterior parietal and medial parietal parcels, with additional high values in occipital and central/sensorimotor-adjacent parcels. Shallower troughs were more evident in frontal/prefrontal, fronto-opercular and cingulate parcels. Thus, the diagonal layer combined a shared spectral motif with anatomical modulation. These results justify treating diagonal precision as the local spectral architecture of the atlas, while the next section focuses on representative off-diagonal real-precision pathways.

### 3.5 Representative real-precision pathways reveal frequency-specific lifespan motifs

The conserved diagonal architecture did not imply that all precision-matrix components followed the same spectral pattern. We therefore examined selected real off-diagonal precision pathways that served as interpretable anchors within the morphology landscape (Figure 6). These pathways were selected from atlas-wide summaries of effect energy, dominant frequency structure, age-shape class and anatomical interpretability. We report HCP-MMP parcel labels, with anatomical descriptions provided where the label is not self-explanatory. The goal was to display representative real-precision motifs rather than to enumerate all source pairs.

**Figure 6.**
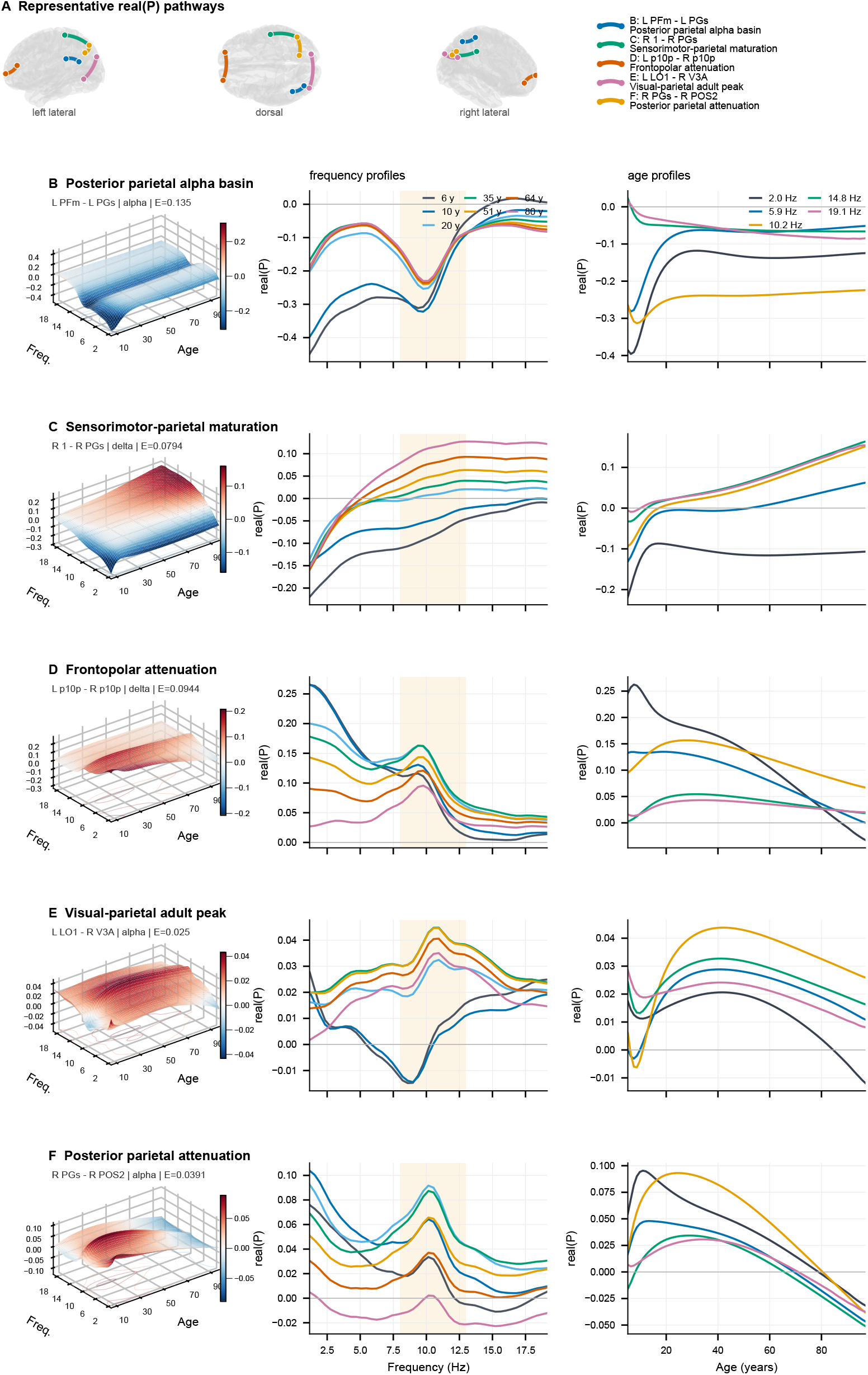
Representative real-precision pathway motifs. **A**, Cortical locations of five selected real normalized precision pathways. **B–F**, Reconstructed real-precision motif cards for posterior parietal alpha basin, sensorimotor-parietal maturation, frontopolar attenuation, visual-parietal adult peak and posterior parietal attenuation. Each row shows the age-frequency surface, frequency profiles sampled at representative ages, and age profiles sampled at representative frequencies. Shaded regions in frequency-profile panels mark the alpha range. HCP-MMP parcel labels are used for pathway names. Pathways were selected as interpretable anchors from atlas-wide screening summaries of effect energy, frequency structure, age-shape class and anatomical location.

The left PFm–left PGs pathway showed a posterior parietal alpha basin (Figure 6B). Its frequency profiles were predominantly negative, but the age slices revealed a change in the location of the strongest depression: the youngest profiles were most negative at the lowest retained frequencies, whereas profiles from young adulthood onward showed a deeper alpha-centered basin near 10 Hz. The age profiles showed the same frequency dependence. Low-frequency and alpha slices became less negative from childhood into adulthood and then approached a plateau, whereas the higher-frequency slices near 15 and 19 Hz started near zero or slightly positive and progressively decreased toward weakly negative values in later life.

The right area 1–right PGs pathway showed a sensorimotor-parietal maturation motif (Figure 6C). Across age slices, the surface retained negative low-frequency values but gained positive values in the alpha and higher retained frequencies. The age profiles were not uniformly increasing across the spectrum: the lowest-frequency slice rebounded early and remained negative, whereas theta-to-high-frequency slices increased steadily from childhood into older adulthood. This pathway therefore combined a persistent low-frequency negative component with progressive strengthening at higher frequencies.

The interhemispheric frontopolar pathway between left p10p and right p10p showed a different low-frequency attenuation motif (Figure 6D). Early profiles were dominated by large positive low-frequency real precision, which declined markedly with age. Mid- and high-frequency slices showed smaller adult peaks, but these did not compensate for the loss of the low-frequency component. Thus, the dominant developmental feature was not a global decrease across the entire spectrum, but a selective attenuation of early low-frequency frontopolar coupling.

The visual-parietal pathway between left LO1 and right V3A had the most explicit shape transition (Figure 6E). In childhood, the frequency profile contained an alpha-range trough, with lower values near 9 Hz than at the flanking frequencies. In adulthood, this trough was replaced by a positive alpha peak centered at a slightly higher frequency, around 11 Hz. The age profiles confirmed this transition: low-to-alpha slices showed inverted-U trajectories with maxima around midlife, whereas the highest-frequency slice was largest at the youngest age and declined across the lifespan. This pathway therefore captured a developmental retuning from an early alpha depression to an adult alpha-centered positive coupling motif.

Finally, the right PGs–right POS2 pathway showed posterior parietal attenuation with a frequency-dependent timing of the peak (Figure 6F). Low-frequency slices peaked earlier, around childhood or early adolescence, whereas the alpha and higher-frequency slices peaked later, from young adulthood into mid-adulthood. By older age, all sampled frequency slices declined, and several crossed toward negative real precision. The surface therefore showed both a shift in the frequency location of the strongest positive component and a late-life attenuation of posterior parietal coupling.

Taken together, the selected pathways illustrate how atlas-level age-frequency variation is expressed in anatomically distinct source pairs. The examples show that a representative real-precision surface may differ not only in overall sign or dominant band, but also in the timing of its peak, the frequency location of its trough or ridge, and whether age changes are shared across frequencies. Thus, Figure 6 gives pathway-level examples of the surface motifs summarized in Figure 4.

### 3.6 Delta precision gradients show an S-A-aligned lifespan hierarchy

The morphology landscape identified low-frequency dominance as one of the major continuous axes of off-diagonal precision variation, and several representative pathways showed their strongest age dependence in low-frequency or delta-range slices (Figures 4 and 6). Motivated by lifespan work describing cortical organization along sensorimotor-association and broader neocortical functional hierarchy axes, we asked whether age-specific delta-band |*P*^⋆^| precision matrices expressed a comparable parcel-level hierarchy [27, 28]. For each target age, we summarized the delta-band precision profile of each parcel with the leading source-precision gradient (Figure 7).

**Figure 7.**
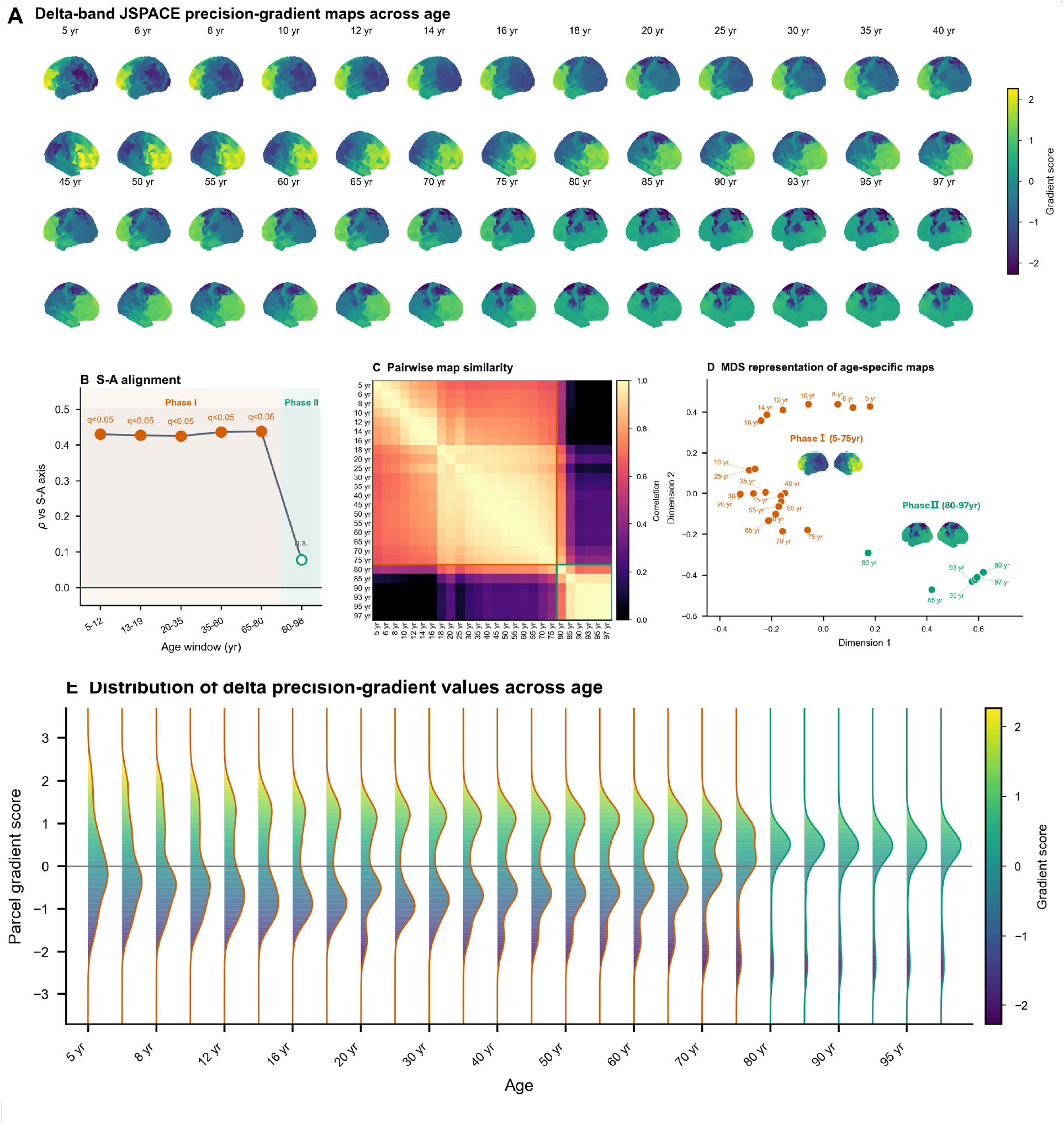
Delta-band JSPACE precision-gradient maps across age. **A**, Age-specific cortical maps of the first delta-band |*P*^⋆^| precision gradient. Values are standardized within each age-specific map. **B**, Alignment between the delta precision gradient and an external sensorimotor-association axis across age windows. Filled markers indicate spin/FDR-supported alignment; the open marker indicates a non-supported oldest-old window. **C**, Pairwise correlation matrix among age-specific gradient maps. **D**, Multidimensional scaling representation of age-specific maps, showing the separation between 5–75 year and 80–97 year map regimes. **E**, Distribution of parcel gradient scores across age. Phase labels denote similarity-defined groups of age-specific precision-gradient maps.

The resulting gradient was aligned with an external sensorimotor-association (S-A) reference from childhood through late life, but this alignment was not observed in the oldest-old window (Figure 7B). Across the 5–80 year windows, correlations with the external S-A axis were moderate and spin/FDR-supported (*ρ* = 0.425–0.438, *q <* 0.05). In contrast, the 80–97 year window contained 19 participants and showed little alignment (*ρ* = 0.078, not supported by the spatial null). Thus, the delta gradient was S-A aligned from 5 to 80 years, whereas the oldest-old maps formed a distinct map regime with weak S-A alignment. The exact transition age should be interpreted with the sparse age support in mind.

The cortical maps showed how this reorganization unfolded spatially (Figure 7A). From childhood through late adulthood, positive gradient scores were concentrated mainly in frontopolar and lateral prefrontal association parcels, including frontoparietal network (FPN)- and default-mode network (DMN)-dominant regions. Negative scores were concentrated in posterior visual-parietal regions early in life and, from young adulthood onward, increasingly in dorsal parietal, cingulate and sensorimotor-adjacent parcels. After 80 years, the positive frontal/prefrontal pole became markedly compressed. By the oldest ages, the highest positive scores were weaker and shifted toward occipital and visuoparietal parcels, whereas the strongest negative scores remained concentrated in parietal-cingulate and sensorimotor-adjacent regions.

Map similarity gave the same transition point. A two-phase description separated maps from 5–75 years and 80–97 years. Within-phase similarity was high for both phases (mean *r* = 0.847 for 5–75 years and *r* = 0.895 for 80–97 years), whereas between-phase similarity was lower (mean *r* = 0.229). The multidimensional scaling (MDS) representation showed older-age maps displaced from the main 5–75 year trajectory (Figure 7C,D).

The distribution plot showed the same late-life shift from another angle (Figure 7E). Because each age-specific gradient was standardized, the vertical spread should be read as the relative ordering of parcels along the gradient, not as a global change in precision magnitude. At 5 years, positive and negative tails were approximately balanced. By young adulthood, the positive tail was more populated, consistent with a prominent association-cortex positive pole. From 80 years onward, no parcels retained strongly positive scores above +1, whereas a negative tail persisted, indicating compression of the positive pole and persistence of a parietal-cingulate negative extreme in advanced age.

## 4 Discussion

JSPACE was developed to estimate frequency-resolved precision matrices of latent cortical sources from scalp cross-spectra. The two simulation benchmarks tested different parts of this goal. The neural-mass benchmark evaluated source-spectral recovery after forward mixing, including power, coherence and phase-lagged structure. The precision-ground-truth benchmark instead evaluated whether a method could recover sparse conditional source relationships. JSPACE performed best for support recovery and leakage-aware localization of true precision edges, whereas amplitude-level partial-correlation recovery remained more difficult. These results support a target-specific conclusion. Source precision is not a substitute for power, covariance or coherence, but a different estimand that requires direct validation when the scientific question concerns conditional source relationships.

A post-hoc source graph can be difficult to interpret when it is fitted to a source moment that has already been shaped by the lead field and inverse regularization. JSPACE reduces this problem by updating posterior source moments and precision matrices within the same forward-model-aware procedure. The standardization step limits the influence of local spectral scale on the graph. The multi-frequency penalty uses neighbouring frequency bins to stabilize estimation. The anatomical weights regularize weakly supported source pairs without fixing the final graph support. Together, these steps make the reported precision less dependent on local power differences, frequency-bin noise and hard anatomical exclusions [1, 17].

In the lifespan atlas, local diagonal terms, off-diagonal source pairs and the delta precision gradient carried complementary information. Diagonal precision showed a conserved alpha-band trough across parcels. Off-diagonal precision surfaces occupied a continuous age-frequency embedding rather than a set of sharply separated edge classes. The leading delta-band precision gradient was moderately aligned with an external sensorimotor-association reference. These levels are linked by one interpretation. JSPACE does not only estimate where source activity is large. It estimates how local and pairwise source relationships remain conditionally structured after other modeled sources have been accounted for.

The diagonal alpha trough illustrates this distinction most clearly. In conventional log-spectrum or covariance surfaces, posterior alpha appears as a peak because alpha has high marginal power. In a precision matrix, the diagonal element has a different meaning. A smaller diagonal precision value indicates larger residual conditional variance for that parcel under the fitted graph. The alpha trough therefore means that alpha-range source fluctuations remain relatively less predicted by the rest of the modeled source system. This does not contradict an alpha power peak. It is the conditional-precision counterpart of a strong local alpha rhythm.

This interpretation is physiologically plausible because resting alpha is partly generated by local posterior and thalamo-cortical loops. Alpha can dominate marginal power while still carrying local timing and gating properties that are not fully explained by other cortical parcels. Work on alpha oscillations supports this view by linking alpha to inhibition, gating and controlled access to information [29, 30]. The posterior and sensorimotor emphasis of the trough depth also fits known alpha and mu rhythm topographies. At the same time, this interpretation should remain component-aware. Alpha-band power is affected by both periodic alpha peaks and aperiodic background activity, and these components can follow different lifespan trajectories [31, 32].

The diagonal surfaces did not separate into distinct region-specific motifs. Instead, all parcels shared a similar alpha-trough pattern, although the depth of the trough varied across systems. Developmental EEG studies help interpret this stability with modulation. In early childhood, the dominant posterior rhythm can lie in theta or low-alpha frequencies. From later childhood into young adulthood, the dominant posterior peak shifts upward within the alpha range [33]. In our atlas, this means that the conserved alpha trough should not be treated as a fixed 10-Hz object throughout life. It is better viewed as an alpha-related conditional feature whose frequency location and depth can shift as posterior rhythms mature and as aging changes the balance between periodic and aperiodic spectral components.

The continuous UMAP embedding gives the off-diagonal counterpart of this result. The source-pair surfaces did not form clean categories because several physiological influences can act on the same pair. Low-frequency attenuation can reflect the reduction of broad slow coordination during maturation. Alpha basins and ridges can reflect posterior rhythm timing. Later high-frequency increases can reflect maturation of long-range synchrony. Late-life weakening can reflect reduced specificity of association systems. These influences are continuous and can overlap. Similar continuum logic appears in lifespan connectome studies, where development is organized along sensorimotor-association axes rather than by isolated regional labels [28, 34, 35].

The posterior alpha-related pathways represent two variants of the same broad motif. PFm– PGs showed a negative basin whose strongest depression moved from lower retained frequencies in childhood toward the alpha range in adulthood. LO1–V3A showed a more explicit retuning, where a childhood alpha-range trough near 9 Hz became an adult positive ridge closer to 11 Hz. A trough-to-ridge transition means that alpha changed from a frequency of relatively weak or negative conditional coupling to the frequency of strongest positive coupling. This pattern suggests a retuning of posterior conditional coordination, not only a rightward shift of an alpha peak. A plausible mechanism is maturation of posterior feedback timing. As conduction becomes faster and more reliable, visual and parietal systems can sustain stable coordination at a higher alpha frequency. This interpretation is consistent with source-level EEG work linking alpha frequency to conduction delay and cortical myelination, and with models showing that delay and coupling jointly shape alpha synchrony [21, 36].

The area 1–PGs pathway showed a different pattern. Its low-frequency component stayed negative or only partially rebounded, whereas theta, alpha and higher retained frequencies became increasingly positive with age. Area 1 is primary somatosensory cortex, while PGs belongs to posterior parietal association cortex. The frequency split suggests that slow somatosensory-parietal fluctuations remained conditionally distinct, while faster components became more positive during maturation. This fits the broader developmental sequence in which primary sensorimotor systems mature earlier and association pathways continue to refine later. EEG evidence also shows that neural synchrony undergoes late restructuring across adolescence, rather than following a simple monotonic increase [34, 37].

The frontopolar p10p–p10p pathway provides the main low-frequency association motif. It showed strong positive low-frequency real precision in childhood and marked attenuation with age. This can be interpreted as a shift away from broad bilateral slow coordination between homologous frontopolar parcels. Frontopolar and lateral prefrontal cortices are late-maturing transmodal systems. During development, increasing segregation of control and default-mode systems may reduce diffuse slow interhemispheric coupling. In later life, these same long-range association systems are vulnerable to disruption of default and control networks [38, 39]. The p10p–p10p surface therefore supports a bounded interpretation. Low-frequency frontopolar conditional coupling is strong early in life, becomes less dominant with maturation and may be vulnerable to later association-system aging.

Posterior parietal and medial parieto-occipital pathways bridge the alpha and low-frequency motifs. PGs–POS2 showed earlier low-frequency peaks and later alpha or higher-frequency peaks, followed by broad decline in older age. This timing suggests that posterior association systems may first rely on slow coordination and later express more frequency-specific feedback or integrative coupling. PGs and POS2 sit near visual, attentional and default-mode interfaces, where lifespan connectivity studies have reported distinct trajectories for precuneus and posterior cingulate systems [40, 41]. The important point is not that this single pathway proves a general rule. Rather, it gives an interpretable example of how low-frequency and alpha-range components can peak at different ages within the same posterior association system.

The delta-band precision hierarchy places these low-frequency motifs at the system level. We compared it with the sensorimotor-association axis because that axis is a compact reference for cortical hierarchy. It separates primary sensory and motor systems from transmodal frontoparietal and default-mode systems [28,42]. The moderate alignment indicates that low-frequency conditional source relationships are partly ordered along this hierarchy. The positive pole in frontopolar and lateral pre-frontal parcels suggests a transmodal association component. The negative pole in posterior, parietal, cingulate and sensorimotor-adjacent parcels suggests a distinct low-frequency profile in posterior and body-related systems. The alignment is therefore meaningful, but incomplete alignment is expected because JSPACE estimates conditional spectral relationships rather than BOLD functional similarity.

The oldest-old deviation should be interpreted as a candidate late-life reorganization. After 80 years, the frontal positive pole became compressed, the strongest positive scores shifted toward occipital and visuoparietal parcels and the negative pole remained prominent over parietal-cingulate and sensorimotor-adjacent cortex. A plausible mechanism is reduced timing specificity in long-range association systems. Myelin and white-matter integrity influence conduction speed and the reliability of inter-areal timing. When these properties decline, long-range frontal loops may become less able to maintain a distinct slow conditional profile. Posterior and sensorimotor-adjacent systems may then contribute more strongly to the leading low-frequency axis. Lifespan diffusion and myelin studies support this mechanism by showing prolonged white-matter maturation followed by age-related decline, but the sparse oldest-old sample means that the exact transition age remains uncertain [43, 44].

The atlas also extends the normative logic of qEEGt and HarMNqEEG from scalp spectral summaries to conditional source relationships [18, 19]. In a clinical setting, a precision-based chart would ask whether an individual’s age- and site-adjusted source interactions deviate from the expected conditional-dependence structure, not only whether local spectral amplitude is abnormal. This could be useful in future studies where altered coordination, network segregation or indirect coupling is more informative than regional power alone. The present study does not test diagnostic classification, but it provides a precision-domain reference for evaluating such applications in clinical cohorts.

Several boundaries constrain these interpretations. The atlas was estimated from 19-channel EEG, so source precision depends on the forward model, parcellation, residual sensor model and regularization. The cohort is cross-sectional, so age effects cannot be equated with within-person developmental trajectories. The representative pathways were selected as interpretable anchors, not as an exhaustive edgewise significance set. The sign of real precision should also be interpreted as a model-derived conditional relation, not as direct excitation or inhibition. These constraints define the appropriate use of the atlas. It is a harmonized source-precision reference that should be tested further with high-density EEG or MEG, longitudinal elderly sampling and multimodal structural or myelin-sensitive imaging.

Taken together, the findings support source precision as a useful coordinate for lifespan EEG connectomics. JSPACE links source inverse modeling with conditional spectral graphs. The resulting atlas suggests a layered organization of lifespan EEG, with conserved alpha-related diagonal structure, frequency-specific off-diagonal motifs and a delta-band hierarchy that partly follows the sensorimotor-association organization of cortex. This precision view complements power, covariance and coherence analyses by asking how age and frequency shape conditional relationships among cortical generators after forward mixing and indirect dependence have been modeled.

## A Notation and symbols

The main symbols used in the estimator, developmental atlas and simulation metrics are listed in Tables A.1, A.2 and A.3. Superscript ⋆ denotes a final selected or refitted quantity in the subject-level estimator and a harmonized reconstructed quantity in the developmental atlas, as specified by the accompanying base symbol.

**Table A.1.**
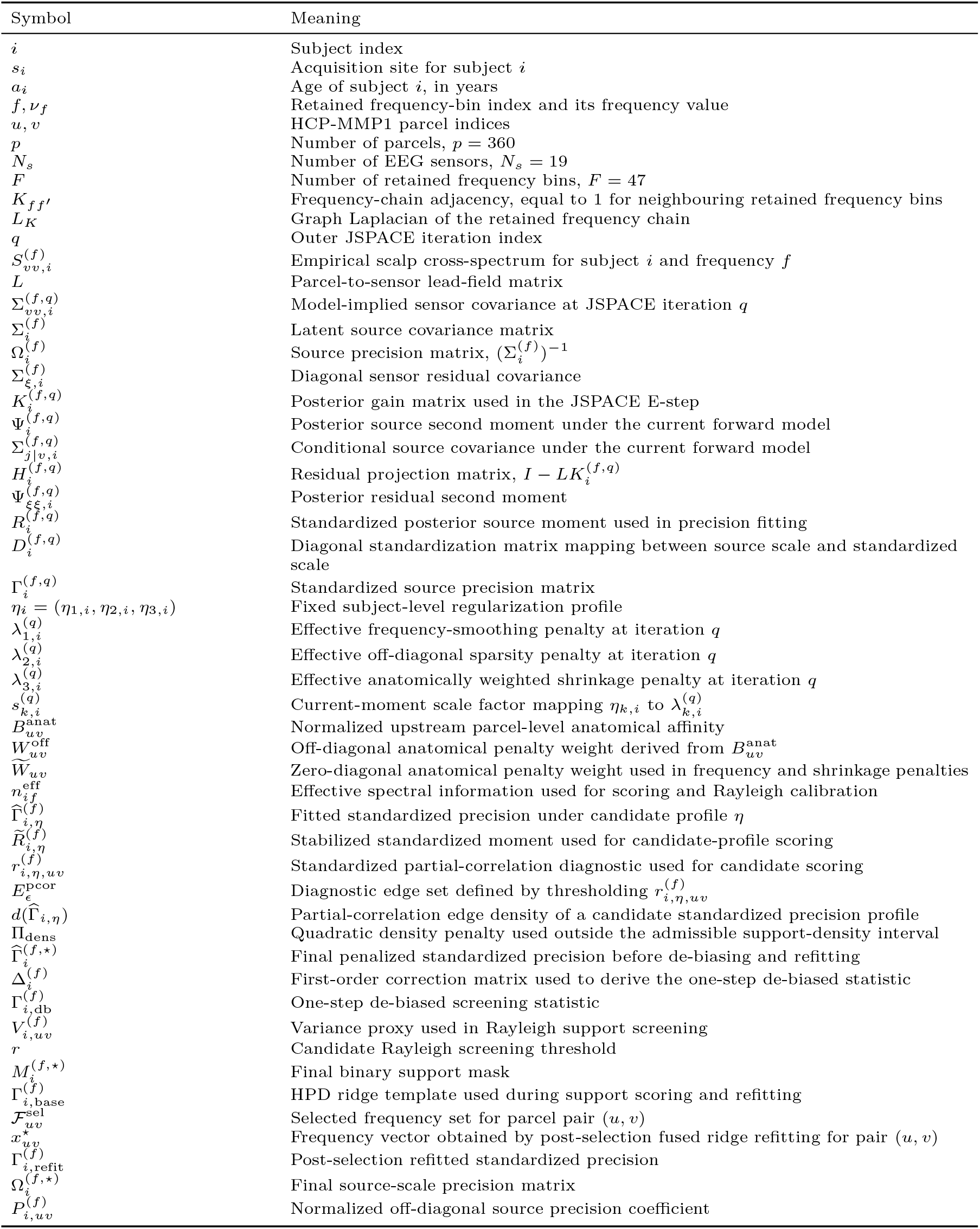
Notation used for the subject-level JSPACE estimator and post-selection procedure.

**Table A.2.**
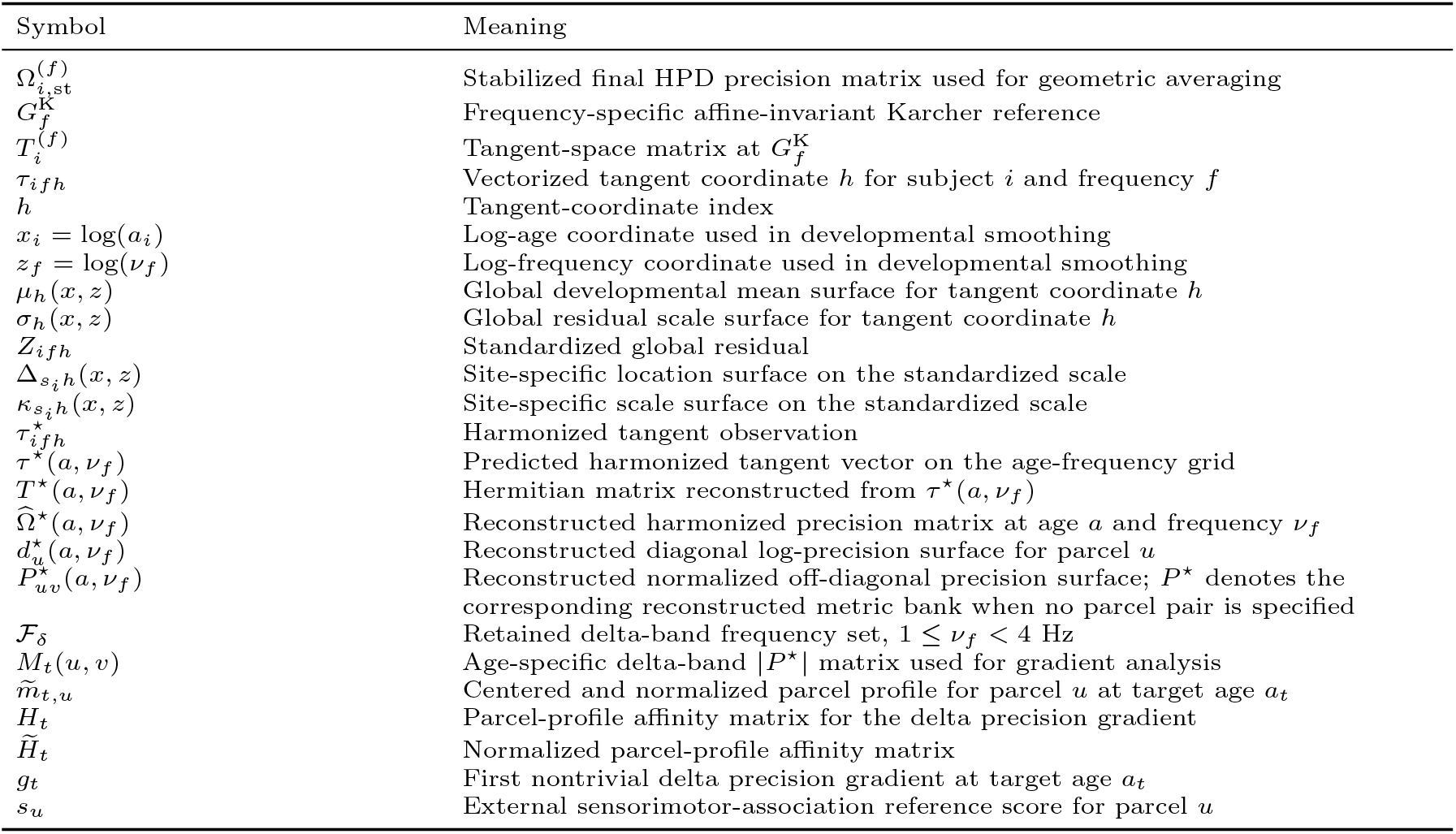
Notation used for the reconstructed Karcher tangent developmental atlas and atlas-wide figure summaries.

**Table A.3.** Notation used only for the simulation benchmark metrics in Figure 3. Symbols in this table are not reused for subject-level JSPACE outputs or developmental atlas surfaces.

| Symbol | Meaning |
| --- | --- |
| $\Omega_{\text{gt}}^{(f)}$ | Ground-truth source precision matrix in the reduced-dimensional precision benchmark |
| $\hat{\Omega}_{\mathcal{A}}^{(f)}$ | Precision matrix estimated by benchmark workflow $\mathcal{A}$ |
| $P_{\text{gt}}^{(f)}$ | Ground-truth normalized partial-correlation matrix derived from $\Omega_{\text{gt}}^{(f)}$ |
| $\hat{P}_{\mathcal{A}}^{(f)}$ | Estimated normalized partial-correlation matrix derived from $\hat{\Omega}_{\mathcal{A}}^{(f)}$ |
| $M_{\text{gt}}^{(f)}$ | Ground-truth off-diagonal support mask at frequency $f$ |
| $\hat{M}_{\mathcal{A}}^{(f)}$ | Estimated off-diagonal support mask for workflow $\mathcal{A}$ at frequency $f$ |
| $M_{\vee}(u, v)$ | Edge-collapsed support mask, defined by the logical OR across frequencies |
| $C_{\text{LF}}$ | Lead-field confusability matrix computed from normalized lead-field columns |
| $C_{\text{PSF/CTF}}$ | Point-spread/cross-talk confusability matrix computed from an MNE inverse response |
| $\mathcal{N}_k^{\text{LF}}(u)$ | Top-$k$ lead-field-confusable neighborhood of parcel $u$ |
| $\mathcal{N}_k^{\text{PSF/CTF}}(u)$ | Top-$k$ PSF/CTF-confusable neighborhood of parcel $u$ |
| $D_{\text{leak}}(u, v)$ | Normalized leakage-derived parcel distance used for spatial edge-localization metrics |
| $d_E(e, \hat{e})$ | Endpoint-swap-invariant distance between a true edge $e$ and estimated edge $\hat{e}$ |
| $TP, FP, FN$ | True-positive, false-positive and false-negative support counts used in $F_1$ and recall metrics |

## B Implementation provenance and cohort tables

The empirical JSPACE analysis used a versioned MATLAB implementation with fixed configuration files. The method takes as fixed inputs a parcel-level EEG forward operator, an upstream parcel-level anatomical affinity matrix, and HarMNqEEG scalp cross-spectral tensors. It does not regenerate individual head models, repeat CiftiStorm–Brainstorm source modeling, or re-run HarMNqEEG pre-processing.

Cohort summaries used for the Methods figures were produced from the retained-subject manifest used in the downstream analysis. Table B.1 reports the retained sample size and age coverage for each recording site.

**Table B.1.** Retained HarMNqEEG cohort composition by recording site. Ages are in years.

| Dataset site | Country | Retained $N$ | Median age | Age range |
| --- | --- | --- | --- | --- |
| New York 1970s | USA | 376 | 12.23 | 5.98–80.40 |
| Chongqing 2016 | China | 256 | 20.00 | 15.00–26.00 |
| Russia 2013 actiChamp | Russia | 231 | 21.00 | 16.00–82.00 |
| Zurich 2017 | Switzerland | 210 | 71.65 | 18.00–89.42 |
| Germany 2013 | Germany | 203 | 27.50 | 22.50–77.50 |
| Cuba 1990 | Cuba | 198 | 18.79 | 5.35–97.00 |
| CHBMP 2004–2008 | Cuba | 133 | 31.01 | 17.50–62.09 |
| Barbados 1978 | Barbados | 70 | 9.24 | 5.18–51.17 |
| Russia 2013 nvx136 | Russia | 68 | 22.50 | 16.00–56.00 |
| Cuba 2003 | Cuba | 55 | 25.84 | 5.17–69.64 |
| Bern 1980 | Switzerland | 46 | 12.96 | 10.17–16.25 |
| Chengdu 2014 | China | 40 | 24.00 | 21.00–28.00 |
| Malaysia 2017 | Malaysia | 28 | 26.00 | 19.00–60.00 |
| Colombia 2019 | Colombia | 21 | 34.00 | 22.00–45.00 |
| Total | – | 1935 | – | 5.17–97.00 |

The reconstructed developmental atlas used the same retained cohort and final JSPACE precision outputs. Table B.2 summarizes the main atlas dimensions and the interpretation layer used in the Results. The site-harmonized *z*-score outputs were retained as harmonization diagnostics, whereas the reported physiological summaries were extracted after reconstruction of the harmonized tangent surfaces back to HPD precision matrices. Cortical surface displays used the fixed Brainstorm/HCP-MMP1 source-space object used by the JSPACE analysis; scout labels and centroids were taken from the same parcel ordering as the 360-node precision matrices.

**Table B.2.** Provenance and dimensions of the reconstructed Karcher tangent developmental atlas.

| Atlas component | Specification |
| --- | --- |
| Participants | 1935 retained HarMNqEEG participants with valid age and JSPACE precision outputs |
| Age grid | 80 target ages spanning 5.17–97.00 years |
| Frequency grid | 47 retained frequency bins spanning 1.1719–19.1406 Hz |
| Parcellation | 360 HCP-MMP1 parcels in the fixed source-space ordering used by JSPACE |
| Tangent representation | 129,600 real components per frequency: 360 diagonal, 64,620 real off-diagonal and 64,620 imaginary off-diagonal tangent coordinates |
| Reconstructed metrics | 360 diagonal log-precision surfaces and 64,620 surfaces for each off-diagonal metric: real normalized precision, imaginary normalized precision, magnitude and phase |
| Primary interpretation layer | Reconstructed harmonized precision metrics $\hat{\Omega}^*(a, \nu)$ and $P_{uv}^*(a, \nu)$ |
| Diagnostic and screening layers | Global and site-harmonized standardized residuals, descriptive surface-screening summaries and source-data tables |

## C Detailed JSPACE estimator

This appendix gives the estimator details that are compressed in the main Methods. For frequency-domain input construction, the empirical scalp cross-spectrum and the effective spectral count were

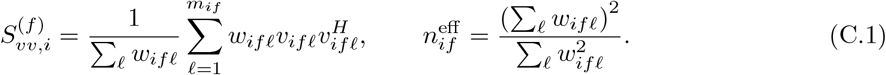

Here *v*_*ifℓ*_ is a sensor Fourier coefficient vector, *m*_*if*_ is the number of retained Fourier coefficients, and *w*_*ifℓ*_ is the corresponding epoch weight. Equal epoch weights give 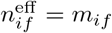.

Structural information entered only through off-diagonal parcel-pair penalties. After symmetrizing the upstream parcel-level anatomical affinity matrix and normalizing it to 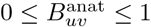, the bounded anatomical penalty weight was

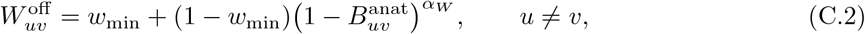

with 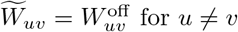 and 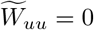.

At outer iteration *q*, the posterior source and residual moments were computed from

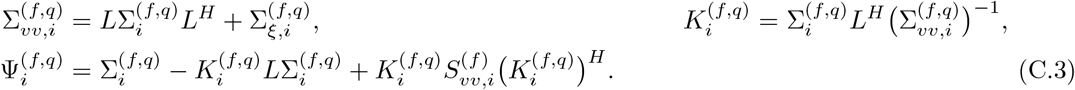

Let 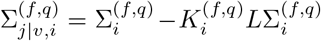 and 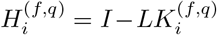. The residual second moment and diagonal residual update were

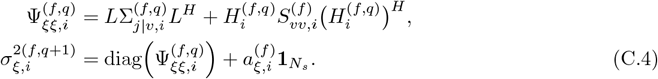

The posterior moment was standardized by

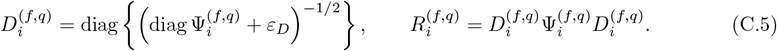

For the frequency-chain adjacency *K*_*ff*_*′* = **1 {**|*f* − *f*^*′*^| = 1 and its graph Laplacian *L*_*K*_, the weighted frequency and shrinkage penalties were

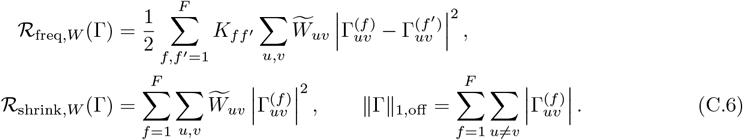

The detailed objective equivalent to the compact main-text objective was

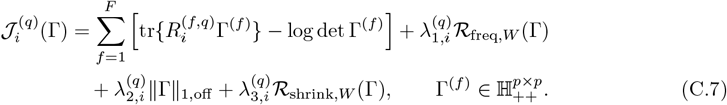

The subject-level regularization profile *η*_*i*_ = (*η*_1,*i*_, *η*_2,*i*_, *η*_3,*i*_) was selected once per subject. Its effective penalties changed across outer iterations through current-moment scale factors:

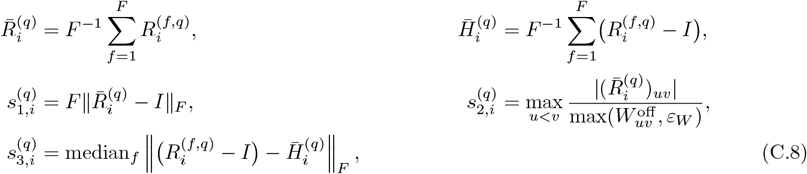

and

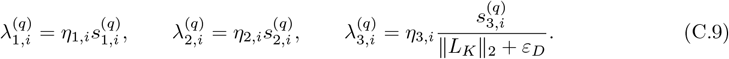

Thus *η*_*i*_ is fixed for a subject, but the penalties used in Eq. (C.7) adapt to the scale of the current standardized posterior moment.

Candidate *η* profiles were ranked by a standardized Gaussian fit, edge-count penalty and density calibration term:

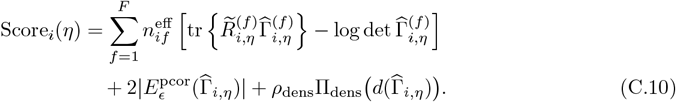

Here 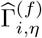 is the fitted standardized precision under candidate profile 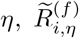 is the corresponding stabilized standardized moment used for scoring, and 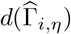 is the partial-correlation edge density. The edge set was computed from standardized partial-correlation diagnostics,

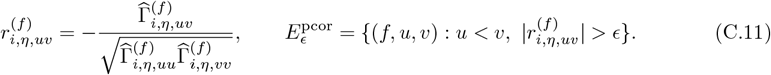

The density penalty Π_dens_ is zero inside the admissible density interval and quadratic outside it; numerical settings are listed in Appendix D.

### One-step de-biasing and Rayleigh support screening

The unpenalized standardized fitting term for one frequency is

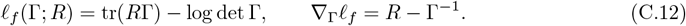

The frequency, sparsity and anatomical terms in Eq. (C.7) act on the standardized source-precision matrices Γ^(*f*)^. The anatomical term uses the parcel-level affinity weights as soft off-diagonal shrinkage and frequency-smoothing weights, not as a hard support constraint. These penalties stabilize the inverse problem, but the off-diagonal terms of the penalized precision are also shrunk toward zero, as in sparse inverse-covariance estimation [15].

Let 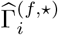 denote the final penalized standardized precision and let 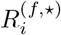 denote the corresponding final standardized posterior moment. Because the penalized fit does not generally satisfy 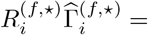 I, JSPACE used a one-step linearized score correction before support selection. Writing 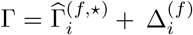 and linearizing the score equation around 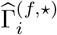 gives

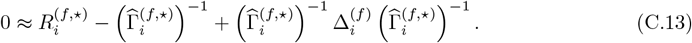

Multiplying this equation on the left and right by 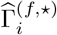 yields

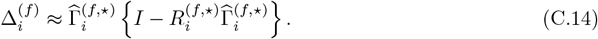

The resulting one-step statistic is

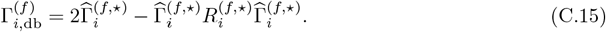

This expression is the inverse-covariance analogue of a one-step de-biased estimator; here it is used only to construct a dense screening statistic, not as the final reported precision matrix [45, 46]. The matrix 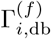 was Hermitianized before thresholding.

Under the null of no edge, the asymptotic linearization treats the real and imaginary parts of an off-diagonal Hermitian precision entry as approximately centered Gaussian components. Under the corresponding circular approximation, the modulus follows a Rayleigh form, consistent with the complex Gaussian and complex-Wishart treatment of spectral matrices [47, 48]. For each candidate Rayleigh threshold *r*, off-diagonal entries were retained when

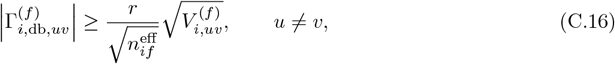

where the variance proxy was computed from the penalized estimate,

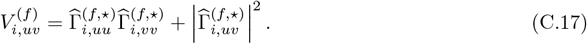

The diagonal entries of a Hermitian positive-definite precision matrix are positive real values; the modulus is therefore needed only for the complex off-diagonal entry. The Rayleigh masks are support-selection devices and were not interpreted as familywise or false-discovery-rate controlled edgewise tests. Diagonal entries were always retained. Candidate Rayleigh masks were scored after applying the mask to the HPD ridge template 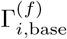, using the same trace-minus-log-determinant support score with frequency, sparsity, anatomical and density penalties. If the selected Rayleigh mask fell outside the admissible density interval, a fallback support was constructed from the final standardized posterior moment to keep the refit numerically stable.

After this support screening, selected off-diagonal entries were refit with a frequency-smoothed ridge problem. Let 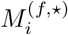 denote the final binary support mask. If 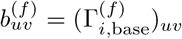 and 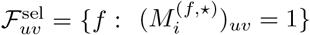, then

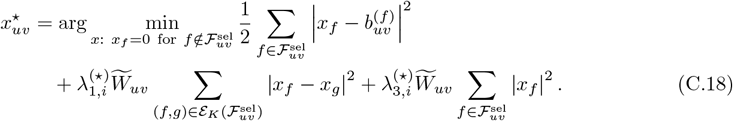

Here 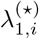 and 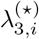 are the final selected effective frequency and shrinkage penalties. The refitted standardized precision was Hermitianized, projected to the HPD cone when necessary, and mapped _back by 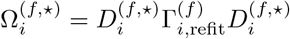._

## D JSPACE implementation constants

The empirical source precision fits used *N*_*s*_ = 19, *p* = 360, and *F* = 47. Effective spectral information in the upstream cross-spectral construction was represented by 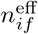, defined in Eq. (C.1). In the HarMNqEEG cross-spectral construction, Fourier coefficients are averaged with equal epoch weights; therefore 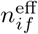 equals the retained epoch count for subject *i* and is constant across the retained frequency bins. This quantity calibrates the hyperparameter scoring and Rayleigh-screening scores, but it does not multiply the normalized stochastic M-step objective.

The empirical fits used 10 outer E/M iterations. The diagonal residual variance vector was initialized separately at each retained frequency as 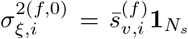, where 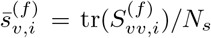. The residual-variance floor in Eq. (C.4) was 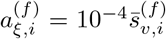, and the update rate was 1.0, so each E-step residual estimate replaced the previous diagonal residual variance before the next E-step. eLORETA initialization used regularization 0.05. JSPACE used the fixed HCP-MMP1 parcel-level anatomical affinity matrix from the upstream structural model used in *ξ*-*α*NET [21]. In that preprocessing, the Rosen–Halgren anatomical connectivity prior was aligned to HCP-MMP1 parcel labels, exponentiated from the stored log-affinity matrix, zero-diagonalized and Frobenius-normalized. JSPACE used this matrix as a continuous soft anatomical penalty weight. The anatomical weight transform used the bounded power form in Eq. (C.2), with *α*_*W*_ = 1.0 and *w*_min_ = 0.2. These values make the anatomical term a soft modulation of the off-diagonal quadratic and frequency-smoothing penalties: *α*_*W*_ = 1.0 gives a linear conversion from normalized anatomical affinity to penalty discount, whereas *w*_min_ = 0.2 bounds the strongest anatomical discount so that no parcel pair is exempt from regularization. A separate high-affinity auxiliary mask derived from the strongest normalized affinities was used only to keep structurally strong pairs eligible in the active-set M-step, not to define the final graph support.

The hyperparameter search used MATLAB `surrogateopt` with 80 maximum function evaluations, 10 minimum surrogate points, no inner parallelism, and common random numbers with base seed 0. The global bounds, fixed after preliminary calibration runs, were

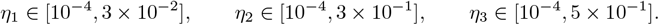

These intervals were obtained from preliminary sweeps across representative subjects and reduced simulation cases to bracket numerically stable HPD behavior, convergence, and plausible sparsity regimes. They were fixed before the final cohort analysis and were not optimized using downstream age, site, or developmental effect surfaces. The initial log-space bank contained

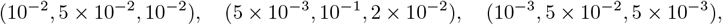

with Latin-hypercube points added to reach 10 initial points. Local search proposals were restricted to ± 0.7 log10 decades around the current search center when this restriction remained inside the global preliminary-calibration domain.

The stochastic M-step used band-stratified sampling with frequency-band edges [0, 4, 8, 13, 20] Hz, nearest-neighbor closure radius 1, backtracking factor 2, maximum 20 backtracking steps, *L*_min_ = 10^−3^, and *L*_max_ = 10^6^. Surrogate objective evaluations used 10 stochastic iterations, whereas the full fixed-point M-step used 30 stochastic iterations and three sampled frequencies per band before nearest-neighbor closure. For local backtracking, let *B*_*r*_ denote the frequency batch at stochastic iteration *r*, let 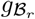 be the smooth part of Eq. (C.7) restricted to the batch and its retained nearest-neighbor terms, let *Y* ^(*r*)^ be the current iterate, let *X* be the candidate update, and let Δ = *X* − *Y* ^(*r*)^. The step was accepted when

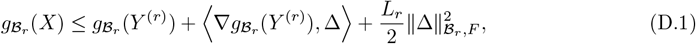

where 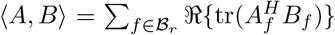 and 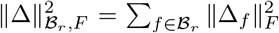. If the inequality failed, *L*_*r*_ was multiplied by the backtracking factor and the proximal step was recomputed until the condition was met or the maximum backtracking count was reached.

The active-mask density floor was max {0.03, log(*p*)*/p*}, equal to 0.03 for *p* = 360. The density lower bound used in tuning and support search was log(*p*)*/p* = 0.01635 for *p* = 360. The upper bound was min(*d*_data_, 0.40), where *d*_data_ denotes the empirical active-mask density before capping, and was then constrained to be at least 1.5 times the lower bound. The density penalty weight was *ρ*_dens_ = 10^7^, the partial-correlation density diagnostic threshold was 3 × 10^−3^, and the support-score edge penalty used *α*_edge_ = 2.

The post-selection Rayleigh grid was 0.7, 0.8, …, 3.1. Candidate hyperparameter fits were ranked by Eq. (C.10); Rayleigh thresholds were ranked by the trace-minus-log-determinant support score in the post-selection section. The post-selection HPD floor was *ε*_post_ = 10^−8^. Let 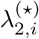 denote the final selected effective *ℓ*_1_ penalty. The ridge template used 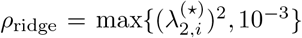 unless an explicit post-ridge penalty was supplied. For this template, if 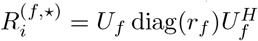, then

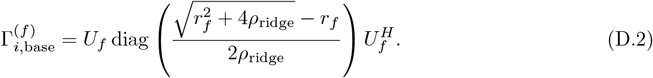

Final precision matrices were obtained with the frequency-smoothed fused ridge refit in Eq. (C.18), followed by condition-stabilized inversion for the paired covariance with target condition number 10^4^ in the scaled precision space.

## E Simulation generation models

The simulation study used two distinct layers: a forward neural-mass covariance benchmark and a reduced-dimensional precision-ground-truth benchmark. The first tests recovery from time-domain source dynamics passed through the EEG forward model. The second tests precision-graph recovery under a known sparse Hermitian precision sequence.

### Forward neural-mass covariance benchmark

For replicate *m*, parcel *u*, and oscillator component *θ* ∈ {*α, ℓ*}, let *x*_*θ,m,u*_(*t*) be displacement and *v*_*θ,m,u*_(*t*) be velocity. The alpha component used *ω*_*α*_ = 2*π*(10 Hz) and *ζ*_*α*_ = 0.05; the low-frequency component used *ω*_*ℓ*_ = 2*π*(3 Hz) and *ζ*_*ℓ*_ = 0.20. Sparse coupling generated the common drive

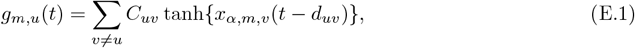

where *C* is a sparse coupling matrix and *d*_*uv*_ are propagation delays. The simulated dynamics were

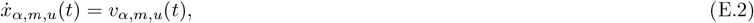

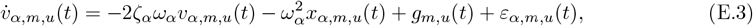

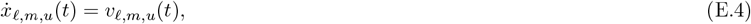

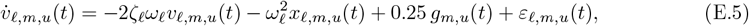

with independent Gaussian source innovations of standard deviation 0.8. The equations were integrated by Euler steps with Δ*t* = 1 ms for 20 s, discarding the first 5 s as burn-in. The source signal was

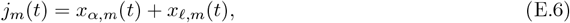

after global source-scale normalization.

Sensor time series were generated by

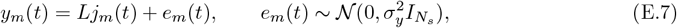

with *σ*_*y*_ set to a fixed fraction of the standard deviation of *Lj*_*m*_(*t*). After burn-in removal, Hann-windowed Fourier coefficients were computed at the target frequencies,

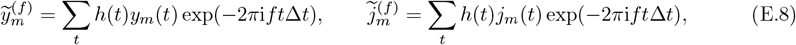

and cross-spectra were ensemble averages

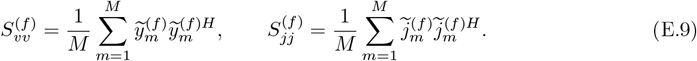

The matrices were Hermitianized and given a small relative ridge. This benchmark does not impose a source-level Wishart perturbation; sampling variability arises from the simulated oscillator ensembles and their Fourier-domain averaging.

The default neural-mass benchmark used 50 replicates, *N*_*s*_ = 8, *p* = 16, six frequencies over 2–20 Hz, *M* = 20 ensembles, sparse coupling density 0.02, coupling strength 0.25, sensor noise level 0.05, and relative ridge 10^−8^.

### Forward benchmark evaluation metrics

Let 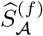 denote the source cross-spectrum estimated by workflow *A* and let 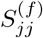 denote the simulated source cross-spectrum. For each frequency, source power was the real diagonal element

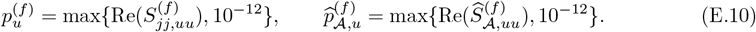

When a workflow returned a fitted log-power surface, the power reconstruction metric compared that fitted surface with the true log_10_ source-power surface:

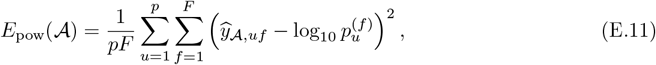

where 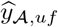 is the workflow-specific fitted log_10_ power value. The direct log-power error used the source covariance diagonal itself,

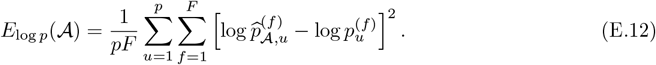

Source-covariance recovery was evaluated by the relative Frobenius error

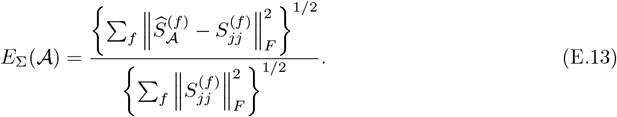

Complex coherency matrices were formed as

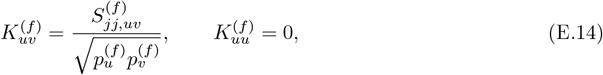

with the same normalization for 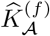. Coherence error used the relative Frobenius error between the complex coherency matrices, and imaginary-coherence error used the same error after replacing each matrix by | Im *K*^(*f*)^|. Peak-frequency error was

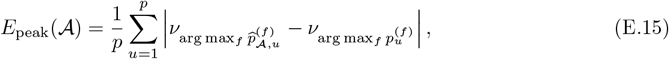

where *ν*_*f*_ is the frequency in Hz. All forward-benchmark metrics were computed for each replicate and then summarized as median and interquartile range in Table 2.

### Reduced-dimensional precision-ground-truth benchmark

The precision benchmark first generated a sparse Hermitian positive-definite source precision sequence. For candidate edge *e* = (*u, v*), an edge envelope was

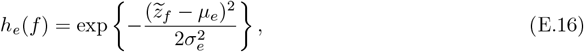

where 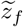 is the normalized frequency coordinate used only for this simulation benchmark. If frequency-varying support was enabled, the edge was active at frequency *f* only when *h*_*e*_(*f*) exceeded the envelope threshold. Conditional on activity,

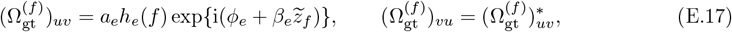

with amplitudes *a*_*e*_ drawn from the specified edge-strength interval and phases drawn independently. Diagonal dominance was imposed by

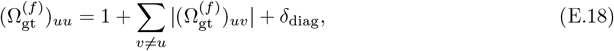

followed, when needed, by a small HPD projection. The source covariance truth was

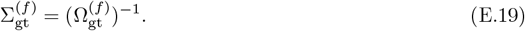

Sensor-level covariance was generated through the same forward model,

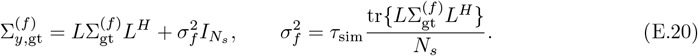

When complex Wishart sampling was enabled, observed sensor spectra were sampled as

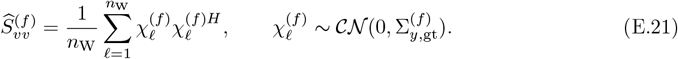

where *n*_W_ is the complex-Wishart draw count. Without Wishart sampling, 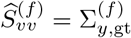. This benchmark provides direct ground truth for precision error, partial-correlation error, support ROC/PR, density bias, and Rayleigh-screening calibration.

The reduced-dimensional precision benchmark used 20 replicates, the real 19-sensor lead-field representation, *p* = 120 selected parcels, 47 frequencies from 1.17 to 19.14 Hz, *n*_W_ = 300, signal-to-noise ratio 13 dB, lead-field perturbation 0.05, target support density 0.08, edge-strength range [0.04, 0.12], diagonal margin 0.40, frequency-varying support enabled, support-envelope threshold 0.25, anatomical-affinity-weighted support sampling when structural affinity was available, zero-lag edge ratio 0.50, and sensor-level complex Wishart sampling enabled.

### Comparator workflows and support extraction

Source-workflow baselines first estimated source covariance matrices independently at each frequency. MNE used the local minimum-norm inverse with regularization 0.05 when the external mn cross routine was unavailable; eLORETA used the corresponding local eLORETA covariance reconstruction. MNE+FOOOF and eLORETA+*ξ*-*α* then fitted source-power spectra for power-level summaries, but precision-level metrics were computed by HPD-stabilizing and inverting the estimated source covariance. Their support masks were defined by thresholding the absolute normalized partial-correlation matrix at 0.10.

The *ξ*-*α*NET baseline used the available structural forward model and source-covariance workflow. The run used 47 frequency bins and the settings

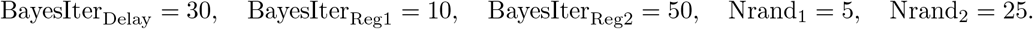

Its source covariance was projected to the HPD cone, inverted and thresholded at the same 0.10 source-workflow partial-correlation threshold.

For MNE+GLASSO and eLORETA+GLASSO, the source covariance was standardized, shrunk toward the identity with covariance-shrinkage weight 0.10, and fitted with an EBIC-tuned Hermitian graphical lasso. The lambda grid contained eight values from *λ*_max_ to 0.15*λ*_max_, with EBIC *γ* = 1.0, ADMM *ρ* = 1, maximum 80 iterations and tolerance 10^−4^. Support was taken from the graphical-lasso support mask when available, otherwise from a precision-entry threshold of 10^−4^ relative to the diagonal scale.

Fused GLASSO used the MNE source covariance by default. It first fit the same EBIC graphical lasso independently by frequency, then applied one-dimensional fused-lasso denoising to each precision entry across frequency. The fusion grid was *λ*_2_ ∈ {0, 0.05, 0.10, 0.20}, selected by total EBIC, with 60 maximum fusion iterations and tolerance 10^−4^.

HIGGS was run frequency-wise with the copied HIGGS baseline using lasso regularization, 100 spectral segments, 60 outer iterations, 30 inner iterations, *α*_*ξ*_ = 10^−4^, no prewarming and a Rayleigh-threshold grid from 0.10 to 1.00 in steps of 0.10. HIGGS covariance estimates were inverted for precision-level metrics.

### Precision-benchmark evaluation metrics

The precision benchmark included exact support metrics and source-imaging-aware spatial metrics. Let 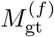 and 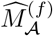 denote the true and estimated off-diagonal support masks for workflow *A* at frequency *f*. Exact support metrics vectorized the upper-triangular off-diagonal entries of all 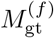 and 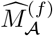 and computed

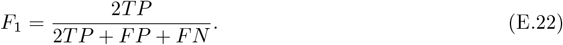

Edge-collapsed support first combined support across frequencies,

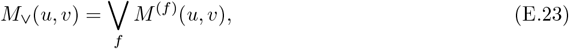

and then computed the same binary support metrics at the parcel-pair level.

Normalized partial-correlation matrices were derived from precision matrices as

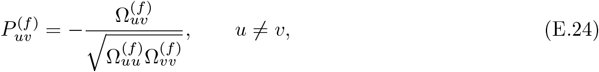

with zero diagonal. The partial-correlation error plotted in Figure 3H was the relative Frobenius error

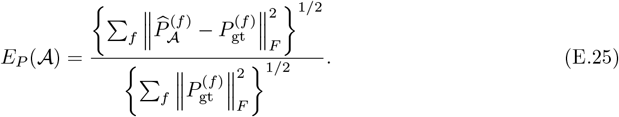

Spatially tolerant support metrics allowed a true edge to match an estimated edge when their endpoints fell into source-imaging ambiguity neighborhoods. For lead-field tolerant metrics, the lead-field columns were first normalized and the confusability matrix was

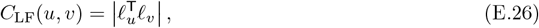

where *ℓ*_*u*_ is the unit-norm lead-field column for parcel *u*. The top-*k* off-diagonal entries of row *u* of *C*_LF_ defined 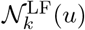, with *k* = 3 and *k* = 5 used in Figure 3C,D.

For PSF/CTF tolerant metrics, an MNE inverse response was formed as

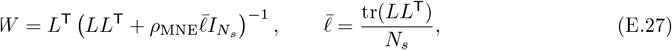

with *ρ*_MNE_ = 0.05. The response matrix *R* = |*WL*| was normalized by both row and column sums and symmetrized to define *C*_PSF*/*CTF_; the top five entries of each row defined 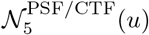.

For either neighborhood definition *N*, a true edge *e* = (*u, v*) matched an estimated edge 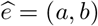 when

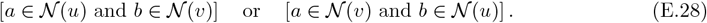

One-to-one matching was used so that a single estimated edge could not explain multiple true edges. Tolerant recall was then *T P/*(*T P* + *FN*) after this spatial matching.

For the distance-based localization metrics, a parcel distance matrix was derived from the leakage/confusability structure. When no explicit simulation distance was available, the distance was

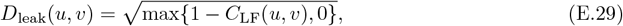

scaled by its non-zero median. The distance between a true edge *e* = (*u, v*) and estimated edge 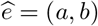 allowed endpoint swapping,

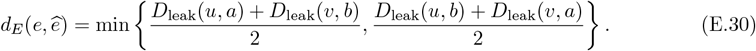

The true-to-estimated edge distance averaged the nearest-estimated-edge distance over true edges. The estimated-to-true distance reversed this direction, and the symmetric edge distance was the mean of the two.

## F Developmental model specification

This appendix gives the equations and implementation settings for the reconstructed Karcher tangent developmental atlas. Let *i* index subjects, *s*_*i*_ acquisition site, *a*_*i*_ age in years, and *ν*_*f*_ the retained frequency. The input to this stage was the final JSPACE precision matrix 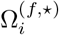.

### HPD stabilization and tangent coordinates

Each matrix was first Hermitianized and stabilized before geometric analysis. In implementation, the matrix was projected to the HPD cone when needed, eigenvalues were floored, and a trace-scaled shrinkage term was applied,

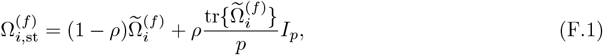

where 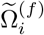 denotes the stabilized HPD projection of 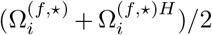. The main run used nearest-HPD projection, an absolute eigenvalue floor of 10^−8^, relative eigenvalue floor 10^−10^, target condition-number control, and shrinkage *ρ* = 10^−4^.

For each frequency, the Karcher reference 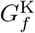 was estimated under the affine-invariant metric. Tangent matrices were

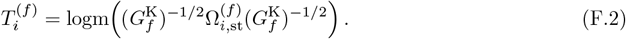

The vectorization map was

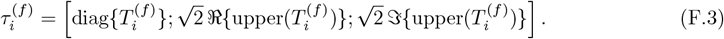

For *p* = 360, this gives 129,600 real tangent components per frequency.

### Global and site-harmonized tangent surfaces

Age and frequency entered the smoother on the transformed coordinates

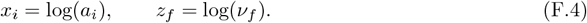

For tangent component *h*, the global mean surface *µ*_*h*_(*x, z*) was fit by bivariate fast local polynomial regression using the fast local-polynomial regression implementation described in FKreg, which accelerates multivariate kernel and local-polynomial regression with non-uniform fast Fourier transforms [49]. The smoother used a Gaussian kernel and local quadratic order. The same engine estimated the residual scale surface *σ*_*h*_(*x, z*) after numerical flooring. The main run used mean-surface bandwidths (0.48, 0.40) and variance-surface bandwidths (0.72, 0.60) on the (*x, z*) axes.

The standardized global residual was

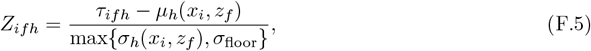

with *σ*_floor_ = 10^−6^. For each site, location and scale surfaces Δ_*sh*_(*x, z*) and *κ*_*sh*_(*x, z*) were estimated on *Z*_*ifh*_ with the same local-polynomial family and bandwidths (0.72, 0.60). Sites with insufficient observations used the global standardized scale rather than an unstable site-specific surface. The harmonized residual was

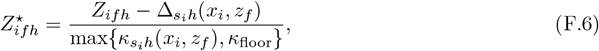

where *κ*_floor_ = 0.25. The harmonized tangent observation was then

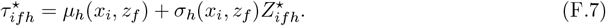

The final atlas surface for component *h* was the local-polynomial mean of 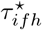 on the target grid. The grid contained 80 ages from 5.17 to 97.00 years and the 47 retained frequencies from 1.1719 to 19.1406 Hz.

### Reconstruction and metric extraction

At target point (*a, ν*_*f*_), the predicted tangent vector *τ*^⋆^(*a, ν*_*f*_) was converted back to a Hermitian matrix *T* ^⋆^(*a, ν*_*f*_) and reconstructed as

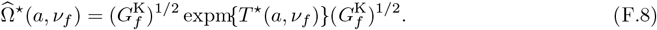

Diagonal and off-diagonal metrics were then

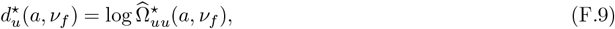

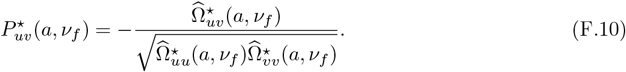

The off-diagonal metric bank stored ℜ (*P* ^⋆^), ℑ(*P*^⋆^), |*P*^⋆^|, and arg(*P*^⋆^) for all 64,620 unique parcel pairs.

### Surface screening summaries

Surface screening summaries were deterministic descriptive summaries of the reconstructed metric surfaces. For age-shape classification, the algorithm computed age-window means, age marginal range, endpoint change, late-life slope, inverted-U score, U-shape score and adolescent-trough score after scale normalization. A surface was classified as adolescent trough when the adolescent-trough score exceeded 0.20, inverted U or U shaped when the corresponding score exceeded 0.18, lifespan increase when normalized endpoint change exceeded 0.20 or normalized late-life slope exceeded 0.01, and lifespan decrease when the corresponding values were below −0.20 or −0.01; otherwise it was classified as weak or flat.

Frequency summaries used the mean absolute value within delta, theta, alpha and beta ranges over the retained grid. The dominant band was the band with the largest mean absolute value. A surface was labelled alpha ridge when the alpha-ridge score exceeded 0.20, higher-frequency dominant when the low-high gradient exceeded 0.20, and lower-frequency dominant when the gradient was below −0.20; otherwise the label followed the dominant band.

Age-frequency interaction strength was computed by comparing the full surface with its additive age-marginal and frequency-marginal approximation. Interaction strength ≥ 0.45 was labelled strong, values ≥ 0.25 and < 0.45 were labelled moderate, and lower values were labelled mostly additive. These summaries were used for atlas screening and representative-panel selection only. They were not treated as edgewise hypothesis tests. Phase summaries were reported as exploratory because phase is angular and the current surface summaries do not yet use circular statistics.

For the continuous morphology landscape in Figure 4, the UMAP input was built from reconstructed off-diagonal |*P*^⋆^| and real *P*^⋆^ surfaces. Each source-pair surface was represented by seven row-centered, root-mean-square normalized feature blocks sampled on 16 age locations and 24 frequency locations: mean-normalized |*P*^⋆^|, log(|*P*^⋆^|+ *ϵ*), the log-magnitude surface after removing the frequency main effect, the age derivative of log magnitude, the frequency derivative of log magnitude, the frequency curvature of log magnitude and signed real *P* ^⋆^. The concatenated feature matrix was feature-standardized, reduced to 80 randomized principal-component analysis (PCA) components, and embedded with UMAP using 60 neighbors, Euclidean distance and fixed random seed 42. The two-dimensional UMAP in Figure 4A was used only as a visualization of surface-shape proximity, not as a clustering result.

The continuous annotation scores in Figure 4 were computed on the full reconstructed grid. Let *u*(*a*) be the frequency-averaged |*P*^⋆^| profile for a source pair and *v*(*ν*) its age-averaged frequency profile. Childhood, late-life and mid-adult windows were defined as *a* ≤ 12, *a* ≥ 60 and ≤ 20 *a* ≤ 40 years. The age-direction score was

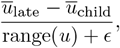

and the midlife-curvature score was

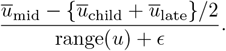

For frequency summaries, delta, theta, alpha and beta bands were defined as 1–4, 4–8, 8–13 and 13–20 Hz within the retained grid. The alpha-axis score was

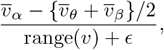

so positive values indicate an alpha ridge and negative values an alpha trough. Low-frequency dominance was

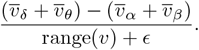

Age-frequency interaction strength was the standard deviation of the residual after subtracting the additive age and frequency main effects from the log-magnitude surface, divided by the standard deviation of the log-magnitude surface. In Figure 4D, feature-extreme sets were defined by the upper or lower 10% of the corresponding continuous score: high age-direction for late increase, low age-direction for late decrease, low alpha-axis for alpha trough, high alpha-axis for alpha ridge, high low-frequency dominance and high interaction strength. Each feature-extreme set therefore contained 6462 of the 64,620 off-diagonal source pairs, and overlaps were used only as a descriptive atlas summary.

### Delta-band precision-gradient and S-A alignment

Figure 7 used a parcel-gradient summary of the reconstructed delta-band precision profiles. For a target age *a*_*t*_, the off-diagonal delta matrix was

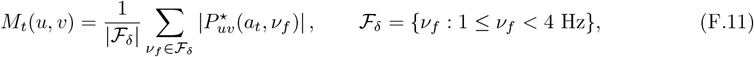

with the diagonal set to zero. Let *m*_*t,u*_ denote row *u* of *M*_*t*_. Each row profile was centered and normalized,

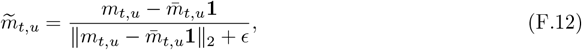

and the parcel-profile affinity was

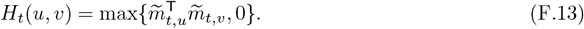

For numerical stability and locality, each row retained its largest *k* affinities,

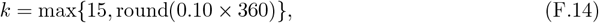

and the matrix was symmetrized. The normalized affinity

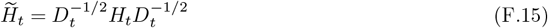

was eigendecomposed, and the first nontrivial eigenvector was used as the target-age precision gradient *g*_*t*_. Each *g*_*t*_ was standardized to zero mean and unit variance across parcels.

The sign of *g*_*t*_ is arbitrary under eigendecomposition. Gradients were therefore oriented to a conservative external sensorimotor-association reference. The external reference was derived from the lifespan S-A gradient reported by Li et al. [27]. Because the external vertex grid and the JSPACE HCP-MMP1 surface are not identical, the main reference used the Li et al. Yeo-7 system gradient mapped to HCP-MMP1 parcels through Schaefer-400 seven-network overlap [50, 51]. For parcel *u*,

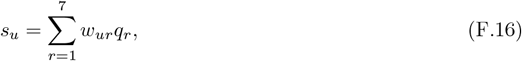

where *w*_*ur*_ is the fraction of the parcel assigned to Yeo network *r* and *q*_*r*_ is the adult S-A score of that Yeo network. The sign of *g*_*t*_ was flipped when its Spearman correlation with *s* was negative.

Panel B of Figure 7 used age-window rather than single-target gradients. The same procedure was applied to matrices averaged over delta-band frequencies and over six age windows: childhood (5–12 years), adolescence (13–19 years), young adulthood (20–35 years), midlife (35–60 years), late life (65– 80 years) and oldest-old (80–97 years). Alignment was quantified by Spearman correlation between the window-specific gradient and *s*. Spatial significance used 5000 within-hemisphere spherical spin permutations of HCP-MMP1 parcel centroids on the Brainstorm registration sphere, with Hungarian assignment used to match rotated positions to parcels. Spin *p* values were two-sided and were FDR-corrected across the band-by-age-window stability grid. Delta-window alignment values and source-subject counts for the corresponding cohort age intervals are reported in Table F.1.

**Table F.1.** Age-window alignment between the delta-band JSPACE precision gradient and the external sensorimotor-association reference used in Figure 7B. Subject counts refer to the source cohort records falling in each age interval. Spatial spin testing used 5000 within-hemisphere rotations, and *q* values are Benjamini–Hochberg FDR corrections across the band-by-age-window stability grid.

| Age window | Subjects | Spearman $\rho$ vs S-A | Spin $p$ | Spin/FDR $q$ |
| --- | --- | --- | --- | --- |
| 5–12 years | 321 | 0.430 | 0.0124 | 0.0469 |
| 13–19 years | 247 | 0.427 | 0.0114 | 0.0469 |
| 20–35 years | 741 | 0.425 | 0.0064 | 0.0469 |
| 35–60 years | 188 | 0.436 | 0.0050 | 0.0469 |
| 65–80 years | 209 | 0.438 | 0.0032 | 0.0469 |
| 80–97 years | 19 | 0.078 | 0.5567 | 0.7494 |

Pairwise map similarity in Figure 7C was computed as the Pearson correlation between ranked target-age gradients. For the MDS and phase display, K-means clustering was applied to the ranked target-age gradients for *k* = 2, …, 8 with 100 random initializations per *k*. The selected two-phase summary corresponded to the maximum silhouette score (0.608). Cluster labels were ordered by median age, yielding a 5–75 year phase and an 80–97 year phase. These phase labels were used only to summarize similarity among age-specific precision-gradient maps.

## Data and Code Availability

The shared raw cross-spectral data with encrypted participant identifiers are hosted on Synapse at https://doi.org/10.7303/syn26712693. Complete access requires a Synapse account and login. The multinational harmonized norms used by HarMNqEEG, including traditional log-spectra and Riemannian cross-spectra, are hosted on Synapse at https://doi.org/10.7303/syn26712979. Code for the JSPACE precision-matrix estimation framework is available at https://github.com/akaKinFish/precision_matrix_estimation. Code for the developmental harmonization and atlas reconstruction analyses is available at https://github.com/akaKinFish/development-jharmn. Generated JSPACE outputs supporting the manuscript figures, including subject-level precision summaries, reconstructed atlas surfaces, figure source tables and plotting-ready source data, will be deposited in a public repository with a persistent identifier before publication.

